# Cryo soft X-ray tomography to explore *Escherichia coli* nucleoid remodelling by Hfq master regulator

**DOI:** 10.1101/2021.11.18.469145

**Authors:** Antoine Cossa, Sylvain Trépout, Frank Wien, Etienne Le Brun, Florian Turbant, Eva Pereiro, Véronique Arluison

## Abstract

Bacterial chromosomic DNA is packed within a membrane-less structure, the nucleoid, thanks to proteins called Nucleoid Associated Proteins (NAPs). The NAP composition of the nucleoid varies during the bacterial life cycle and is growth phase-dependent. Among these NAPs, Hfq is one of the most intriguing as it plays both direct and indirect roles on DNA structure. Indeed, Hfq is best known to mediate post-transcriptional regulation by using small noncoding RNA (sRNA). Although Hfq presence in the nucleoid has been demonstrated for years, its precise role is still unclear. Recently, it has been shown *in vitro* that Hfq belongs to the bridging family of NAPs. Its bridging mechanism relies on the formation of the amyloid-like structure of Hfq C-terminal region. Here, using cryo soft X-ray tomography imaging of native unlabelled cells and using a semi-automatic analysis and segmentation procedure, we show that Hfq significantly remodels the *Escherichia coli* nucleoid, especially during the stationary growth phase. Hfq influences both nucleoid volume and absorbance. Hfq cumulates direct effects and indirect effects due to sRNA-based regulation of other NAPs. Taken together, our findings reveal a new role for this protein in nucleoid remodelling that may serve in response to stress conditions and in adapting to changing environments. This implies that Hfq regulates nucleoid compaction directly via its interaction with DNA, but also at the post-transcriptional level via its interaction with RNA.

## INTRODUCTION

Hfq is a pleiotropic regulator found in about 50% of sequenced Gram-positive and Gram-negative bacteria^1^. While its function in Gram(*+*) is still unclear^2^, it was initially discovered as an *Escherichia coli* host factor required for bacteriophage Qβ RNA replication^3^. Even if *hfq* is not an essential gene, its importance in RNA metabolism and in many related cellular functions was demonstrated by the observation of numerous adverse phenotypes in *hfq*^*-*^ cells^4^. These unfavourable effects of *hfq* mutation are linked to the role of Hfq in the stimulation of RNA annealing, an activity called “RNA chaperone”^5, 6^. In particular, Hfq interacts with <u>s</u>mall non-coding RNAs (<u>s</u>RNAs) and facilitates base-pairing of sRNAs with their target mRNAs^7^. The sRNA:mRNA complexes formed typically inhibit mRNA translation and often lead to subsequent RNA degradation^8^. Less frequently, sRNAs also activate RNA translation^9^. A well-known example of sRNA-based regulation consists of the annealing of MicA, a small noncoding RNA of *E. coli*, to *ompA* mRNA, which encodes an outer membrane porin^10^. The roles of Hfq and sRNA are particularly important for bacterial stress resistance, when bacteria must survive in changing conditions^11, 12^. Indeed, Hfq is involved in bacterial virulence as it helps respond to new conditions encountered in the host during infection^13, 14^. This role of Hfq in virulence is illustrated with a *Vibrio cholerae* Δ*hfq* mutant that is highly attenuated in a mouse model infection^15^. In turn, Hfq and sRNA are also involved in antibiotic resistance^16^, bacterial communication and quorum sensing, processes that could also influence bacterial pathogenicity^17^.

The RNA chaperone function of Hfq is primarily due to its N-terminal region (NTR). Indeed, this region of the protein adopts a typical fold, called Sm-fold^18^. Structurally this fold is characterized by a secondary structure consisting of an α helix followed by a twisted five-stranded β sheet. This β sheet allows the self-assembly of Hfq into its functional form, a ring-shaped homo hexamer^19^. The surface of the Sm ring comprising the N-terminal α-helices is designated as the proximal face; the opposite face is referred to as the distal face. Both surfaces of the NTR have been shown to bind RNAs with different specificities and affinities^19-22^. In *E. coli*, each Hfq monomer consists of 102 amino acid residues in which the Sm/NTR region contains about 65 residues. But in addition to this NTR, Hfq also comprises a C-terminal region (CTR) of about 35 residues. The CTR region extends outside the NTR, at the periphery of the torus^23^ and is preferentially located at the proximal side of the torus^24^. Structurally, this region has been described for years as intrinsically disordered and indeed, to date, its precise 3D structure is still unknown^7, 25, 26^. Recently, a new property has been discovered for this region as the CTR has been shown to be able to convert to an amyloid structure under certain conditions^27, 28^. Notably, the presence of a nucleic acid promotes this conversion from an intrinsically disordered region into an amyloid domain^29^. This transition has been shown to strengthen the hexameric structure^23^. The role of the CTR on RNA annealing is controversial. Nevertheless, it seems to be dispensable for most sRNA-based regulations^23, 30^, even if the lack of this CTR could affect a few sRNA-mediated regulations^31-34^.

Besides these RNA-related functions, Hfq has also been shown to interact with DNA^35^ and is found in the nucleoid, i.e., the membrane-less viscous structure where the bacterial chromosome is compacted by proteins^36^. Hfq thus belongs to the family of Nucleoid-Associated Proteins (NAPs), jointly with a dozen of other DNA-binding proteins^37-39^. Hfq’s role in DNA structuring is unclear but it is the third most prevalent protein in the nucleoid during the exponential growth phase^37^ and its concentration increases by a factor of two during the stationary growth phase^40^. While other NAPs such as H-NS (<u>h</u>istone-like <u>n</u>ucleoid <u>s</u>tructuring protein), HU (<u>h</u>eat-<u>u</u>nstable nucleoid protein), Dps (<u>D</u>NA-binding <u>p</u>rotein from <u>s</u>tarved cells) or IHF (integration <u>h</u>ost factor) seem to be uniformly distributed in the nucleoid, Hfq seems to have a heterogeneous localisation pattern^41^. As its average concentration in the nucleoid is about 10-20% of total Hfq (∼10-15 µM)^37, 40^, its local concentration could thus locally reach tens to hundreds of µM. This high concentration could thus results in the amyloid assembly of Hfq via its CTR^42^.

Usually, NAPs have three DNA-binding modes which are: (*i*) bending (examples are HU, IHF, Dps), (*ii*) wrapping (i.e., leucine-responsive regulatory protein, Lrp) or (*iii*) bridging (H-NS). Hfq belongs to the third family of bridging NAPs and significantly compacts DNA^29, 43-46^. The Hfq bridging property presumably helps the bacterial chromosome (4.6 Mb, 1.5 mm long) to fit *in vivo* within the small bacterial nucleoid volume (∼0.2 μm^3^)^47^. While Hfq-NTR interacts non-specifically with the DNA backbone^25^, the compacting potential relies mainly on the CTR region of Hfq, independently of the Sm/NTR region^45, 48^. Because Hfq has a preference for A-tract repeats^44, 49^, that are found regularly throughout bacterial DNA and may influence promoter efficiency^50, 51^, Hfq is presumably important for DNA packing and the control of genetic expression. Indeed, some phenotypic effects due to the lack of Hfq initially attributed to RNA regulation^4^ may be directly linked to defects in DNA-related processes^4, 16, 52-56^.

While numerous experiments point to an important role for Hfq and in particular of its CTR in DNA packaging, its precise influence on this process *in vivo* still remains poorly understood. To address this question, we used cryo soft X-ray tomography (cryo-SXT). This technique allows DNA imaging in 3D at high resolution (∼ 30 nm half pitch) while preserving bacterial cells in a near-native state^57^. Indeed, in contrast to eukaryotic cells, the lack of membrane surrounding the bacterial DNA makes its observation *in vivo* challenging without staining (for instance by using diamidino-phenylindole DAPI^58^). One major advantage of this technique, compared to high resolution fluorescence techniques working below the diffraction limit (PALM/STORM^59^) is to be able to analyse label-free DNA^60, 61^, as DNA staining may influence its structure and compaction. Our cryo-SXT analysis demonstrates the influence of Hfq on nucleoid compaction in particular during the stationary growth phase of *E. coli*, and to illustrate the role of Hfq-CTR in this process. These results confirm a new mechanism of DNA bridging by a NAP using an amyloid structure *in vivo* and unveil a novel function for Hfq in nucleoid remodelling that adapts to changing environments.

## RESULTS AND DISCUSSION

As Hfq effects on DNA structure may be direct or indirect (see our previous work about the effect of Hfq on DNA topology^29^), we analysed three bacterial strains: (*i*) MG1655 WT (Wild Type, reference strain), (*ii*) MG1655-Δ*hfq* (totally devoid of Hfq protein) and (*iii*) MG1655-Δ*ctr*, expressing a truncated form of the protein with only the first 72 amino-acids corresponding to the NTR^29^. Each strain was studied during exponential and stationary growth phases, generating a total of 6 different conditions. Bacteria were plunge-frozen on electron microscopy grids. Cryo-SXT tilt-series were collected to study the 3D structure of bacteria. A set of representative bacteria (at least 3 bacteria per condition) out of 67 were selected and are described throughout the whole analysis workflow for the sake of clarity and transparency. Note that each voxel in the SXT reconstructed volume corresponds to the linear absorption coefficient (LAC) (referred to as the absorbance in the text) of the material comprised within it^62^. The absorbance, which translates into the density, of the various analysed nucleoids can then be compared. A collection of reconstructed tomogram slices is presented to appreciate the morphological differences between the different strains (Figure 1).

**Figure 1.**
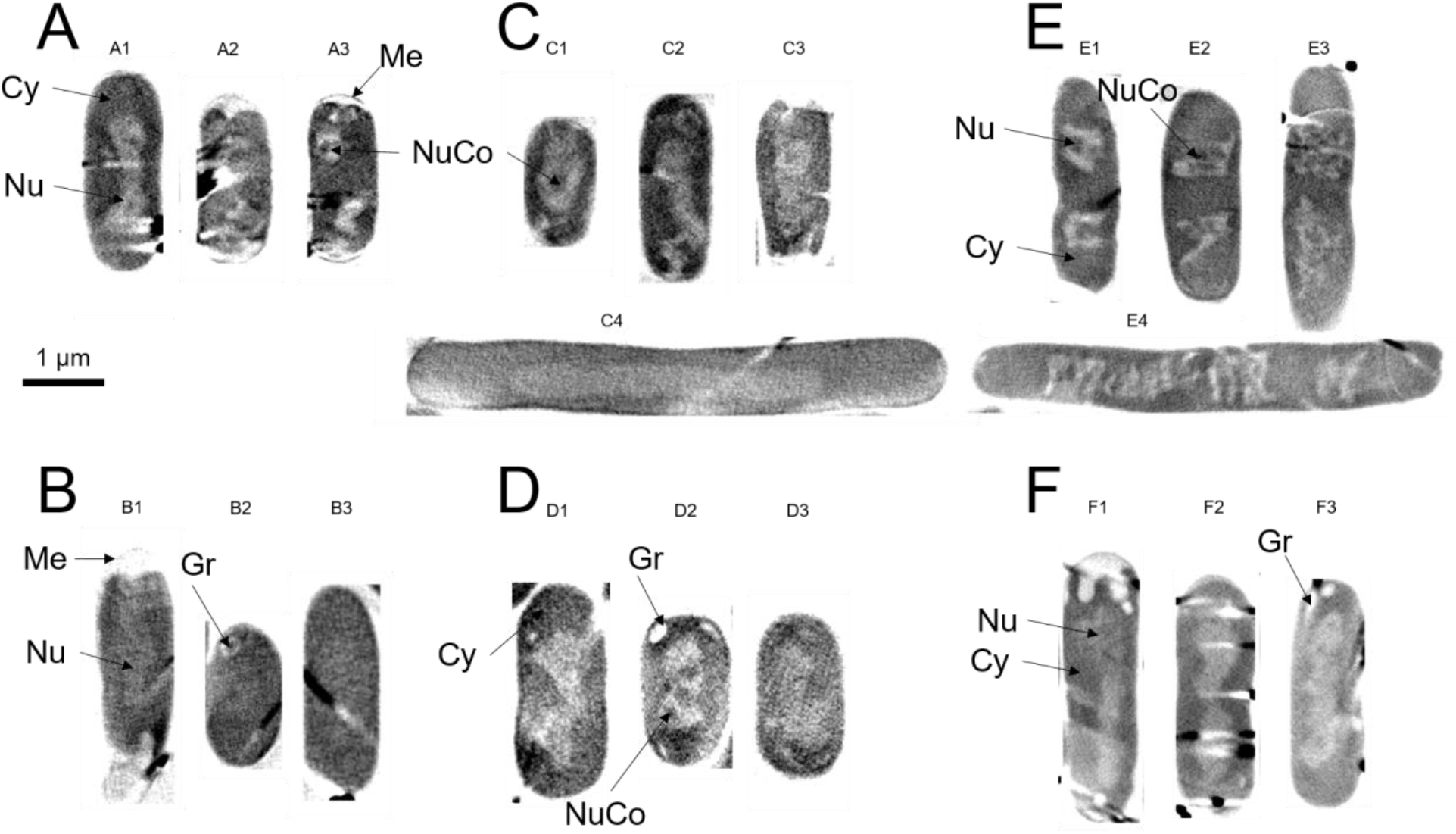
Tomographic reconstructed slices of representative bacteria for the different strains and growth phases. The images have been extracted from absorbance reconstructed volumes which contrast has been inverted to resemble that of transmission images. A) Exponential phase WT cells. B) Stationary phase WT cells. C) Exponential phase Δ*hfq* cells. D) Stationary phase Δ*hfq* cells. E) Exponential phase Δ*ctr* cells. F) Stationary phase Δ*ctr* cells. Scale bar is 1 µm. Several cellular structures have been highlighted: the nucleoid (Nu), the nucleoid core (NuCo), the cytoplasm (Cy), the membrane (Me) and the granules (Gr).

Nucleoids (Figure 1, Nu) can clearly be observed inside the cells as clear zones (lower absorption) inside the darker cytoplasm (higher absorption) (Figure 1, Cy). Depending on the cell growth phase, this nucleoid can be oval-shaped, dumbbell-shaped or round-shaped just after replication. Some nucleoids have a hollow structure (Figure 1, NuCo) that is particularly visible on some bacteria (Figure 1, bacteria A3, C1, D2 and E2). Note that from these various shapes arise heterogeneities in the statistics discussed below. Membranes can also be observed (Figure 1, Me in bacteria A3 and B1), nevertheless due to the resolution of the method (estimated at about 26 nm based on the zone plate used^57^), the double membrane characteristic of Gram-negative bacteria is usually not resolved. In stationary phase cells, granules, possibly polyphosphate granules based on their shape, size and contrast^63-65^, can also be observed (Figure 1, Gr in bacteria B2, D2 and F3). Indeed, polyphosphate granules are expected to appear as low density material (i.e., white pixels) because they possess a high Oxygen content, which generates almost no contrast in the water window in cryo-SXT. Reconstructed tomogram slices of more bacteria are available in Supplementary Figures 1-6. The nucleoid volume, absorbance and shape of the different strains were then analysed and compared. The choice of these three strains allowed us to discriminate indirect effects due to sRNA-based regulations, mainly due to Hfq-NTR, from those directly due to DNA compaction by Hfq-CTR^45, 48^.

### Evolution of nucleoid volume and absorbance between exponential and stationary phases in WT cells

The analysis presented in Figure 2 shows that the nucleoid volume is significantly greater (p-value = 0.0002532) in exponential phase cells (mean = 0.251 µm^3^) compared to stationary phase ones (mean = 0.208 µm^3^). The absorbance of the nucleoid also evolves as it is significantly less dense (p-value = 0.001183) in exponential phase (centred around 0.500 µm^-1^) compared to stationary phase (centred around 0.550 µm^-1^) (Figure 2B and Figure 3A, left panel, red and blues curves respectively). The mean, median and standard deviation values of the different populations are summarised in Table 1.

**Table 1.** Summary of the nucleoid volumes and optical densities mean, median and standard deviation values. These values are associated with the box plots of Figure 2.

| | Volumes ($\mu\text{m}^3$) | | | | LACs ($\mu\text{m}^{-1}$) | | |
| --- | --- | --- | --- | --- | --- | --- | --- |
|  | Mean | Median | Std |  | Mean | Median | Std |
| WT expo | 0.251 | 0.248 | 0.157 | WT expo | 0.493 | 0.505 | 0.0305 |
| WT stat | 0.208 | 0.189 | 0.153 | WT stat | 0.542 | 0.547 | 0.0371 |
| $\Delta hfq$ expo | 0.318 | 0.287 | 0.169 | $\Delta hfq$ expo | 0.496 | 0.495 | 0.0271 |
| $\Delta hfq$ stat | 0.314 | 0.273 | 0.156 | $\Delta hfq$ stat | 0.486 | 0.484 | 0.0231 |
| $\Delta ctr$ expo | 0.357 | 0.359 | 0.109 | $\Delta ctr$ expo | 0.575 | 0.577 | 0.0125 |
| $\Delta ctr$ stat | 0.479 | 0.434 | 0.169 | $\Delta ctr$ stat | 0.603 | 0.612 | 0.0353 |

**Figure 2.**
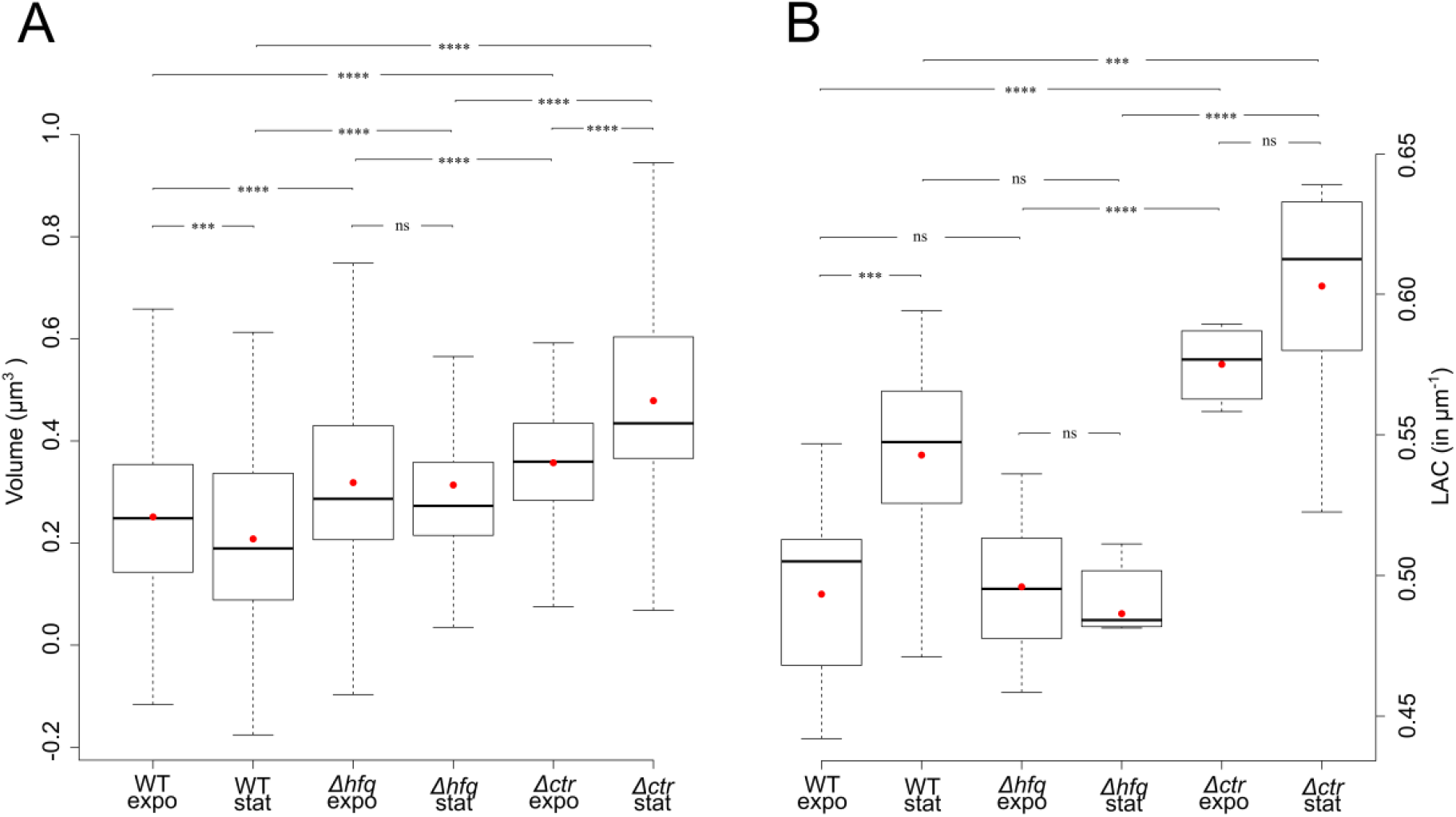
Summary of the nucleoid volumes and optical densities. A) Box plot representation of the nucleoid volume of the different bacteria strains. B) Box plot representation of the mean nucleoid absorbance (LAC) of each bacterium per strain type. The mean and median values are indicated by a red dot and a bold black horizontal line, respectively.

**Figure 3.**
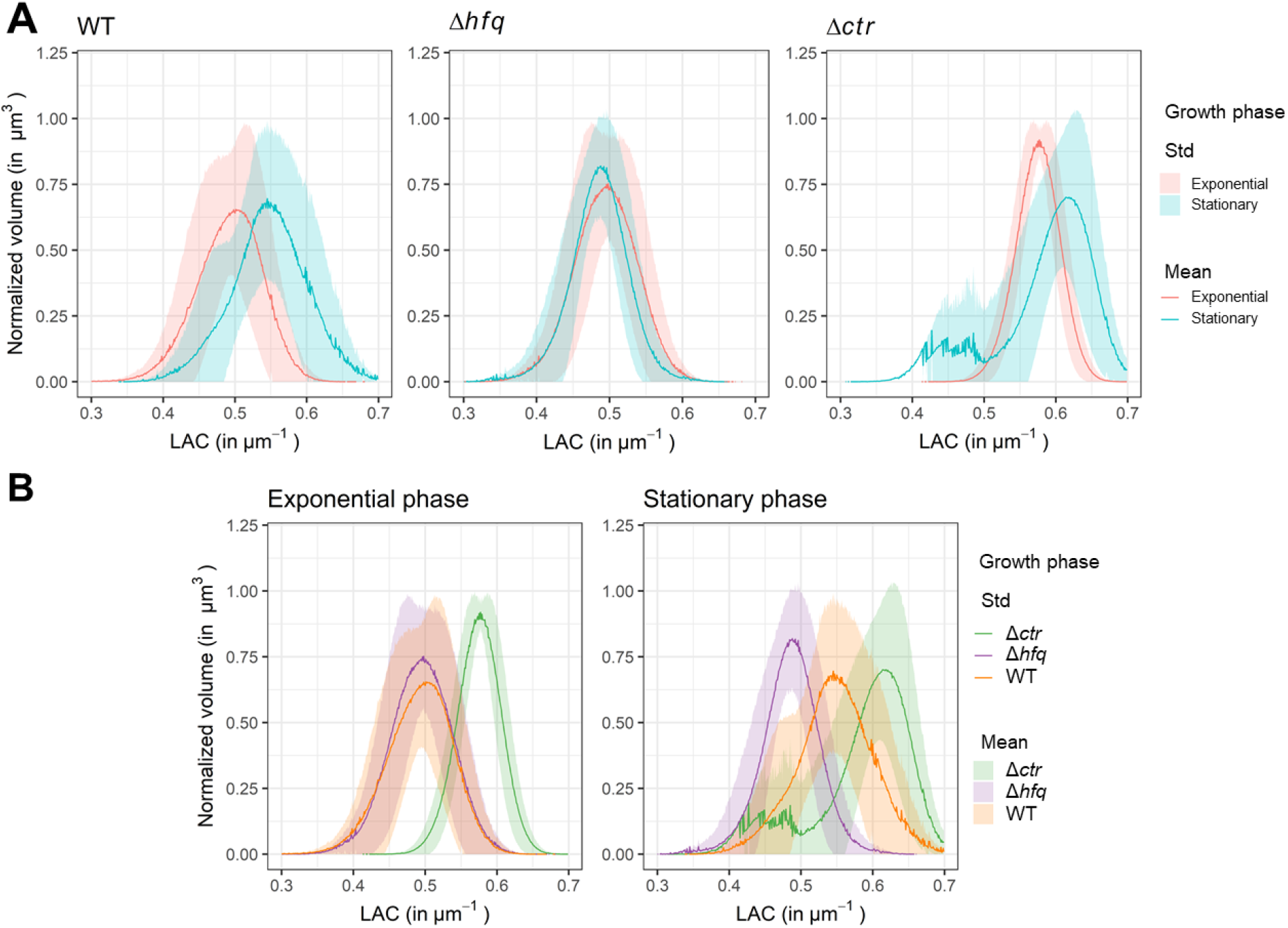
Histograms of nucleoids volumes as a function of LAC for each *E. coli* strain. Results are displayed to highlight both the effects of the growth phase for each strain (A) and the effects of the different *hfq* mutations during each growth phase (B). A) LAC histograms per strain: in red exponential phases and in cyan stationary phases. B) LAC histograms per growth phase: in orange the WT strain, in purple the Δ*hfq* strain and in green the Δ*ctr* one. Histograms represent the mean value (intense line) and the standard deviation (transparent).

Overall, in WT cells, the LAC is shifted to higher values during stationary growth phase, which therefore indicates higher nucleoid absorbance. More precisely, the distribution of the LAC values is bimodal. There is a first sub-population centred around 0.475 µm^-1^ for both growth phases, and there is a second sub-population centred around 0.520 µm^-1^ in the exponential phase which is shifted towards 0.550 µm^-1^ in the stationary phase (Figure 3A, left panel). In both growing conditions, the main sub-population is the one with the highest LAC values. This bimodal distribution could reflect the coexistence of bacteria sub-populations, as they are not synchronized for their division. These results are in agreement with previous studies as the nucleoid absorbance has already been described to be higher for stationary-phase cells compared to exponential phase growing ones^61, 66^. Note that the results presented here do not show a strong reduction of the nucleoid volume as was observed in other studies^61^. This observation may be explained by the fact that the analysis was carried out on mid-stationary phase cells, due to the difficulty of reaching late stationary phase for the Δ*hfq* strain, due to the positive effect of Hfq on σ^S^, the stress and stationary phase sigma factor, expression^67^. As described later in this work, WT cells were compared to *hfq* mutants grown in the same conditions, focusing on mid-stationary phase was mandatory for our analysis. One would have expected a more important nucleoid volume reduction in late stationary phase, as described previously with the same reference strain MG1655^61^. To summarise, in the presence of Hfq the transition from exponential to stationary phase reduces the nucleoid volume while increasing its absorbance. Possibly, the absorbance/density increase observed here is a direct consequence of decreased volume.

### Effect of Hfq deletion on nucleoid volume and absorbance in Δhfq cells

In the Δ*hfq* strain, the nucleoid volumes in exponential phase cells (mean = 0.318 µm^3^) and in stationary phase cells (mean = 0.314 µm^3^) are not statistically different (p-value = 0.6467) (Figure 2). Moreover, in the Δ*hfq* strain, the nucleoid absorbance does not change significatively between exponential and stationary phases (p-value = 0.8048). The LAC values of both growth phases are centred between 0.490 and 0.500 µm^-1^ (Figure 2B and Figure 3A, middle panel). The distribution of the LAC values is bimodal for the exponential phase growth condition (first sub-population centred around 0.475 µm^-1^ and second one centred around 0.510 µm^-1^). In the absence of Hfq, neither the volume nor the absorbance of the nucleoid are significantly modified when the cells shift from exponential to stationary phase.

The comparison with WT cells reinforces this analysis. The nucleoid volumes of exponential phase WT cells (0.251 µm^3^) and exponential phase Δ*hfq* cells (0.318 µm^3^) are statistically different (p-value < 0.0001) (Figure 2) whereas the distributions of their LAC values, again related to nucleoid density, superimpose (LAC centred around 0.500 and 0.490 µm^-1^, respectively and p-value = 0.8656) (Figure 2B and Figure 3B, left panel, orange and purple curves respectively). Interestingly, the small sub-population of the LAC value bimodal distribution in exponential phase WT cells corresponds to the large one in exponential phase Δ*hfq* cells, and *vice versa* (Figure 3B, left panel, orange and purple curves respectively). Thus, in exponential phase, the absence of Hfq influences the volume of the nucleoid, but not its absorbance and density. Moreover, during the stationary phase, the nucleoid volume is significantly greater (p-value < 0.0001) (Figure 2) and the absorbance is not significantly lower (p-value = 0.009324, using Bonferroni correction, see Material & Methods) in Δ*hfq* cells compared to WT cells (Figure 2B and Figure 3B, right panel, orange and purple curves respectively). The LAC value distribution of the stationary phase Δ*hfq* cells are centred at 0.490 µm^-1^ which is about the same position of that of the small sub-population of stationary phase WT cells (Figure 3B, right panel, purple and orange curves respectively). This suggests that stationary phase Δ*hfq* cells could not enter a state that is normally reached by stationary phase WT cells. Taking into account previous *in vitro* analysis, these results were expected, as the absence of Hfq should result in less compaction and thus in an increase of the volume and/or a decrease in absorbance^45, 48^. Here we observe an effect on the volume only, similarly to what has already been observed for the exponential phase condition. Hence, these results strongly suggest a positive role for the Hfq protein on the nucleoid volume, in particular during the stationary phase, when Hfq is more abundant^40^. Note that the major reorganization of *E. coli* nucleoid upon entering the stationary phase was previously attributed to Dps and IHF NAPs^68^. Here we show that Hfq may also play a significant role on nucleoid structure (direct or indirect, see below) during the stationary phase of growth.

### Effect of Hfq-CTR deletion on nucleoid volume and absorbance in Δctr cells

As Hfq effects could be indirect *via* sRNA-based regulations^29^, the outcome of CTR deletion was then tested by studying Δ*ctr* cells. Indeed, Hfq-CTR is dispensable for most sRNA-based regulations, even if few exceptions exist^30, 31^. In short, deletion of the CTR should abolish the direct effect of Hfq on DNA compaction^48^, while most sRNA-regulations controlled by Hfq-NTR should remain^19^. The analysis shows that the nucleoid volume is significantly smaller (p-value < 0.0001) in exponential phase Δ*ctr* cells (mean = 0.357 µm^3^) compared to stationary phase Δ*ctr* cells (mean = 0.479 µm^3^) (Figure 2). The absorbance of the nucleoid is not statistically less dense (p-value = 0.02524) in exponential phase compared to the stationary phase, even though the mean LAC changes from 0.575 to 0.615 µm^-1^ (Figure 2B and Figure 3A, right panel, red and blue curves respectively). The absence of significance is mainly due to the presence of low LAC values in the stationary phase that induces an important skewness on the left part of the curve (Figure 3A, right panel, blue curve). Even though the distribution of the stationary phase Δ*ctr* cells mean LAC values cannot be rejected as being normal (Shapiro normality test p-value = 0.1148), this important skewness could represent a distinct population. Interestingly, this putative distinct population has an average LAC value (inferior to 0.5 µm^-1^) corresponding to that of the small sub-population of stationary phase WT cells, but also to that of the stationary phase Δ*hfq* cells (Figure 3B, right panel, green, orange and purple curves, respectively). In absence of the Hfq-CTR, the transition from exponential phase to stationary phase increases the nucleoid volume but not the nucleoid absorbance.

Compared to WT condition values, we observe that the deletion of the CTR significantly induces an increase of the volume (p-value < 0.0001) and of the absorbance (p-value < 0.0001) during the exponential phase of growth (volume increases from 0.251 to 0.357 µm^3^ and LAC increases from 0.500 to 0.575 µm^-1^). During the stationary phase, the deletion of the CTR also induces a significant increase (p-value < 0.0001) of the volume (from 0.208 to 0.479 µm^3^) and a significant increase (p-value = 0.004096) of the absorbance (LAC increases from 0.550 to 0.615 µm^-1^). Thus, the absence of the CTR domain is associated with an increase of the volume and absorbance during both the exponential and stationary phases compared to WT conditions. As the CTR is expected to compact DNA, its absence should result in a less dense nucleoid occupying a larger volume. This is what is observed for the volume. Conversely, the absorbance/density of the nucleoid is increased.

Compared to the Δ*hfq* cells, the results show that the nucleoid volume and absorbance in Δ*ctr* cells are significantly increased in both growth phases. The volume changes (p-value < 0.0001) from 0.318 to 0.357 µm^3^ and the LAC increases (p-value = 0.0003108) from 0.500 to 0.575 µm^-1^ in exponential phase; and the volume changes (p-value < 0.0001) from 0.314 to 0.479 µm^3^ and the LAC increases (p-value < 0.0001) from 0.490 to 0.615 µm^-1^ in stationary phase. Thus, the deletion of Hfq-CTR in Δ*ctr* cells generates a new phenotype, different from that of the complete deletion of Hfq in Δ*hfq* cells. Several hypotheses could explain these observations:

i. The presence of the Hfq-CTR domain could influence Hfq stability, as shown previously *in vitro*^23^ and thus its abundance *in vivo*. The effect observed could thus be due to different concentrations of full-length Hfq or truncated Hfq. We can rule out this hypothesis as the relative amounts of Hfq and its CTR-truncated form were similar (Supplementary Figure 7). The differences observed thus rely on the regulation mode exerted by Hfq when its CTR is present or not, not on its abundance in the cell.
ii. It is thus likely that the NTR part of the protein, still present in the Δ*ctr* strain, allows a sRNA-regulation that influences nucleoid compaction. A well-known example of sRNA-based regulation by Hfq implies DsrA, a small noncoding RNA of *E. coli*, at several different mRNA targets. These include *rpoS* and *hns* mRNA, which encodes σ^S^, the stress and stationary phase sigma factor^67, 69^ and the nucleoid-structuring protein H-NS^70^, respectively. While DsrA sRNA positively regulates the expression of *rpoS* in particular during the stationary phase, it negatively regulates *hns* mRNAs translation^71^. Note that if Hfq-CTR has recently been shown to be able to regulate *rpoS* expression using DsrA, the Hfq-NTR part alone is sufficient to allow the regulation^34^. One possibility is that Hfq-NTR may regulate H-NS expression. Nevertheless, it was recently shown that H-NS does not significantly affect nucleoid volume and compaction^72^. Thus, Hfq most likely regulates other NAP(s) directly or indirectly. Hfq may for instance regulate positively the expression of another NAP, directly or indirectly if the expression of this NAP is under the control of σ^S^.
iii. One can also imagine that Hfq and precisely its NTR that firmly binds to RNA^19^, could in turn stabilize some noncoding sRNA directly involved in nucleoid structuring^22^, similar to noncoding RNAs shaping of chromatin^73^.
iv. Finally, we cannot rule out that Hfq-NTR may be involved in protein-protein interactions with other NAPs and that these interactions are regulated during the cell cycle^74^. This is particularly important as this regulation might be dependent on the abundance and availability of the proteins, and in particular of Hfq. The Hfq concentration in the nucleoid during the stationary phase is twice that during exponential phase^40^ and in these conditions the protein could be more prone to form amyloid-like structures^27, 28^. The amyloid aggregation of Hfq-CTR could have other indirect effects through potential protein-protein interactions mediated by a specific conformation of the CTR (intrinsically disordered *vs* amyloid). Hfq could thus be differently available for potential interactions with other proteins, a mechanism which would be more important during the stationary phase because of the higher protein concentration. In Δ*ctr* cells, this CTR-mediated control is absent and such an interaction would not occur.

### Nucleoid shape in WT cells

In parallel, we also analysed the spatial distribution of the nucleoid domains as a function of their absorbance. The same bacteria as that previously described in Figure 1 were used. In exponential phase WT cells, low-absorbance nucleoid (Figure 4A, yellow isosurface) and high-absorbance nucleoid (Figure 4A, red isosurface) interdigitate as both isosurfaces are simultaneously visible, which is particularly visible in bacteria A1 and A2. Moreover, hollow regions located deep inside the nucleoid, as already observed in Figure 1, have an absorbance different from that of the nucleoid, and as such, were not detected as part of the nucleoid during the semi-automatic segmentation analysis. Blue spheres have been manually placed to locate the hollow regions in the 3D isosurfaces (Figure 5A). These regions have been described and designated as “core” in previous reported literature^59, 75^. In stationary phase WT cells, the low-absorbance nucleoid (Figure 4B, yellow isosurface) completely surrounds the high-absorbance nucleoid which is consequently barely visible (Figure 4B, red isosurface). Interestingly, the cores or hollow regions previously detected are reduced or disappear during the stationary phase, a dynamic remodelling in agreement with previous reports^59, 75^.

**Figure 4.**
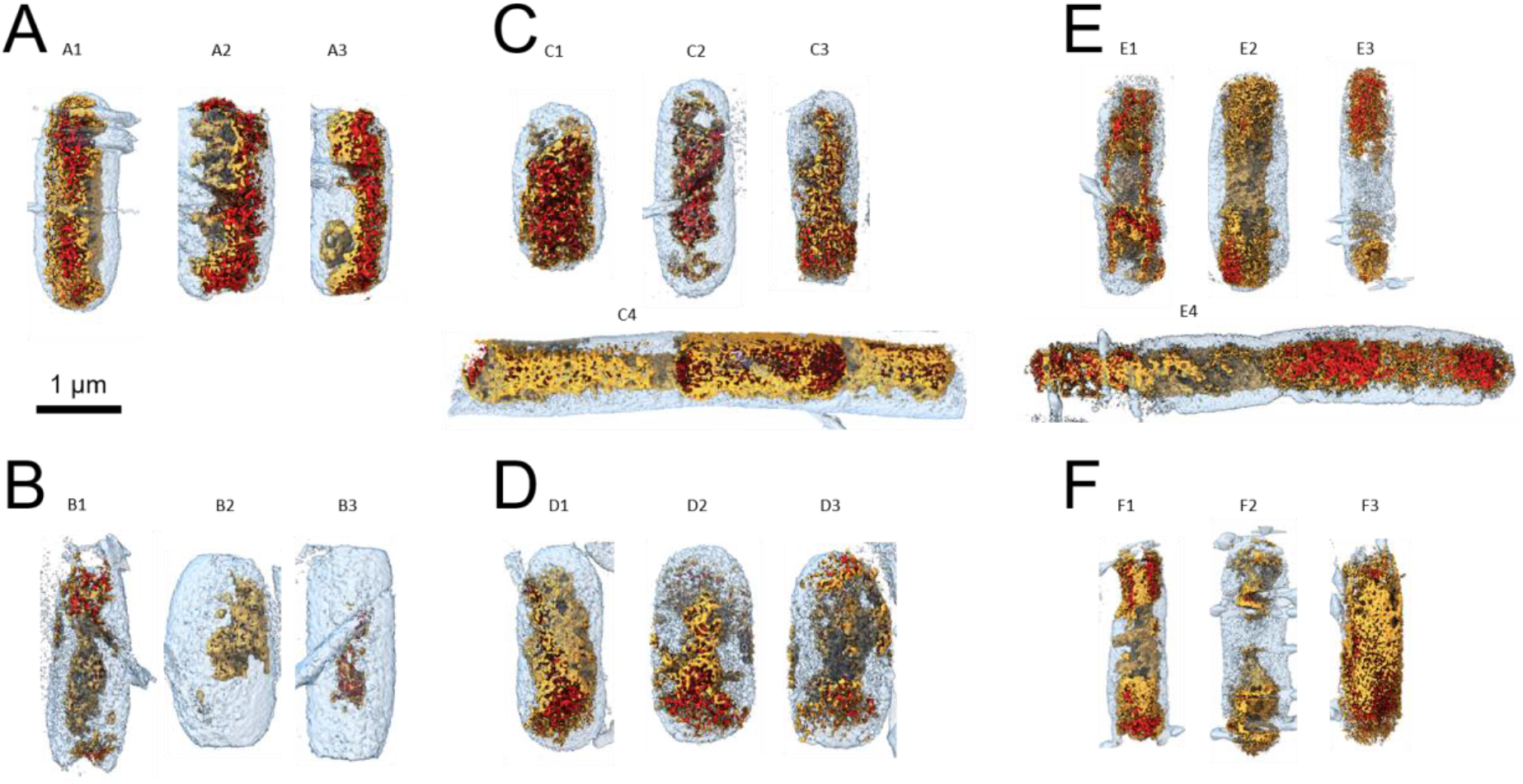
Visualisation of the nucleoid volume and absorbance for each bacteria strain. The different segmentations have been automatically generated using a unique algorithm which was equally applied to all bacteria. A) Exponential phase WT cells. B) Stationary phase WT cells. C) Exponential phase Δ*hfq* cells. D) Stationary phase Δ*hfq* cells. E) Exponential phase Δ*ctr* cells. F) Stationary phase Δ*ctr* cells. The blue isosurfaces are segmentations of the bacteria membranes. The red and yellow isosurfaces are segmentations of the high-absorbance and low-absorbance regions of the nucleoid, respectively. These segmentations allow discrimination of loose nucleoids from compact ones. In loose nucleoids, the high- and low-absorbance nucleoid regions interdigitate, whereas low-absorbance nucleoid surrounds and hides the high-absorbance one in compact nucleoids.

**Figure 5.**
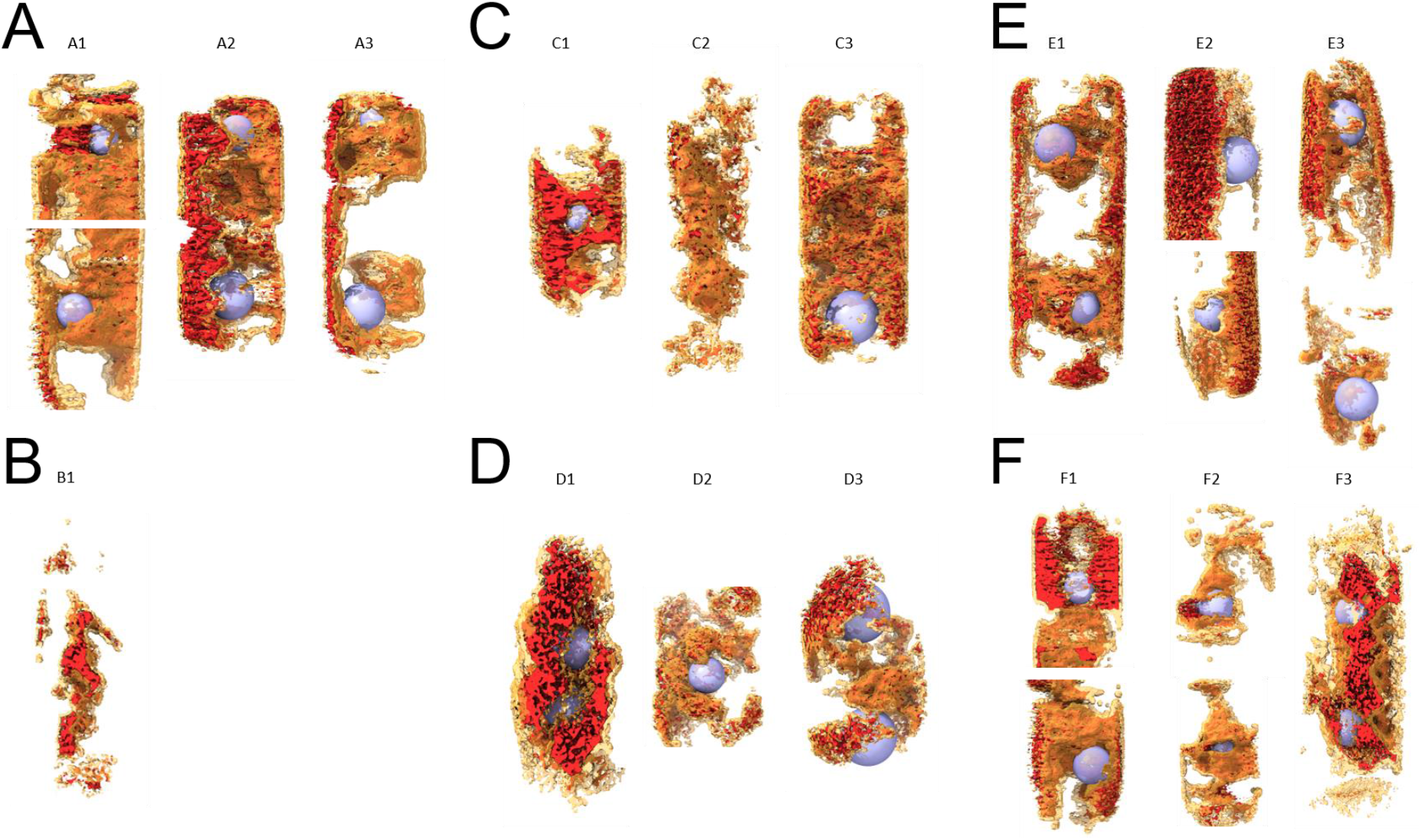
Visualisation of the nucleoid cores. A) Exponential phase WT cells. B) Stationary phase WT cells. C) Exponential phase Δ*hfq* cells. D) Stationary phase Δ*hfq* cells. E) Exponential phase Δ*ctr* cells. F) Stationary phase Δ*ctr* cells. In some of the bacteria, the nucleoid is characterised by the presence of one (or more) hollow region(s) which are known as nucleoid cores, previously observed in tomographic projections of the bacteria (Figure 1). These cores are highlighted using transparent blue spheres which have been manually placed at the centre of the hollow regions. Hollow regions were not observed in stationary phase WT cells.

In stationary phase, the concentration of full length Hfq or its dense packing (following the amyloid aggregation hypothesis) prevents the formation of these core domains. These core domains may presumably be Ori, Ter, Left, and Right arms^76^, even if these domains cannot be formally identified with cryo-SXT only. Correlative SXT imaging in combination with fluorescence and/or cryo-transmission electron microscopy using labelled marker proteins^40, 77^ could help to answer this question.

### Effect of Hfq and its CTR on nucleoid shape

In Δ*hfq* and Δ*ctr* cells, the low-absorbance nucleoid (Figure 4C-F, yellow isosurfaces) and the high-absorbance one (Figure 4C-F, red isosurfaces) interdigitate too, both in exponential phase and stationary phase. Moreover, the presence of cores is also detected in these bacteria, both in exponential and stationary phases (Figure 5). These core domains are similar to those observed in exponential phase WT cells. Thus, it turns out that Hfq and its CTR does not only influence the absorbance and the volume of the nucleoid (DNA compaction) but also its shape. This analysis unveils that the absence of the core domains in stationary phase WT cells is directly or indirectly dependent on the presence of Hfq-CTR. As mentioned above, in stationary phase WT cells (i.e., when the CTR is present and when Hfq concentration is important), the core domains disappear. We now observe that in stationary phase Δ*ctr* cells (i.e., when the CTR is absent and when Hfq concentration is high), the core domains are present. This suggests that the Hfq protein at high concentration, and more particularly its CTR, has a role in the disappearance of the core domains.

### Effect of Hfq and its CTR on cell shape

Finally, elongated cells were observed in the Δ*hfq* strain (Figure 1, bacteria C4). Such elongated cells were not observed in the WT strain. We thus confirm that the absence of Hfq results in the formation of some elongated cells, as previously described^4^. Interestingly, we also observed this phenotype for the strain expressing Hfq devoid of its C-terminal region (Δ*ctr*) (Figure 1, bacteria E4). We then performed the first analysis of the volume, absorbance and shape of nucleoids in *hfq* mutant elongated cells. Based on the nucleoid semi-automatic segmentation, these elongated cells also have interdigitated low-absorbance and high-absorbance nucleoid domains (Figure 4, C4 and E4, bottom bacteria), as in regular-size bacteria. The volume and absorbance of the elongated nucleoids were 1.797 µm^3^ and 2.074 µm^3^ and 0.4882 µm^-1^ and 0.6248 µm^-1^ for these Δ*hfq* and Δ*ctr* cells, respectively.

### Concluding remarks

Taken together our analysis confirms (*i*) that the role of Hfq on DNA structuring exists not only *in vitro* but is confirmed *in vivo* and (*ii*) that its interaction with DNA *via* its CTR induces nucleoid remodelling that may serve for DNA-related regulation (such as replication or transcription) to adapt to changing environments (Figure 6). We show that Hfq amyloid CTR directly influences nucleoid volume and absorbance, but additional effects due to Hfq-NTR related to sRNA-based regulation also exist. These effects are particularly critical during the stationary phase when Hfq is the most abundant (Figure 6B). Thus, our analysis reinforces the role of NAP for this master regulator to answer to stresses encountered by bacteria during their life in adverse environments such as during host colonization^78^, not only by using the sRNA response but also by mediating DNA compaction, thanks to the functional amyloid property of Hfq-CTR.

**Figure 6.**
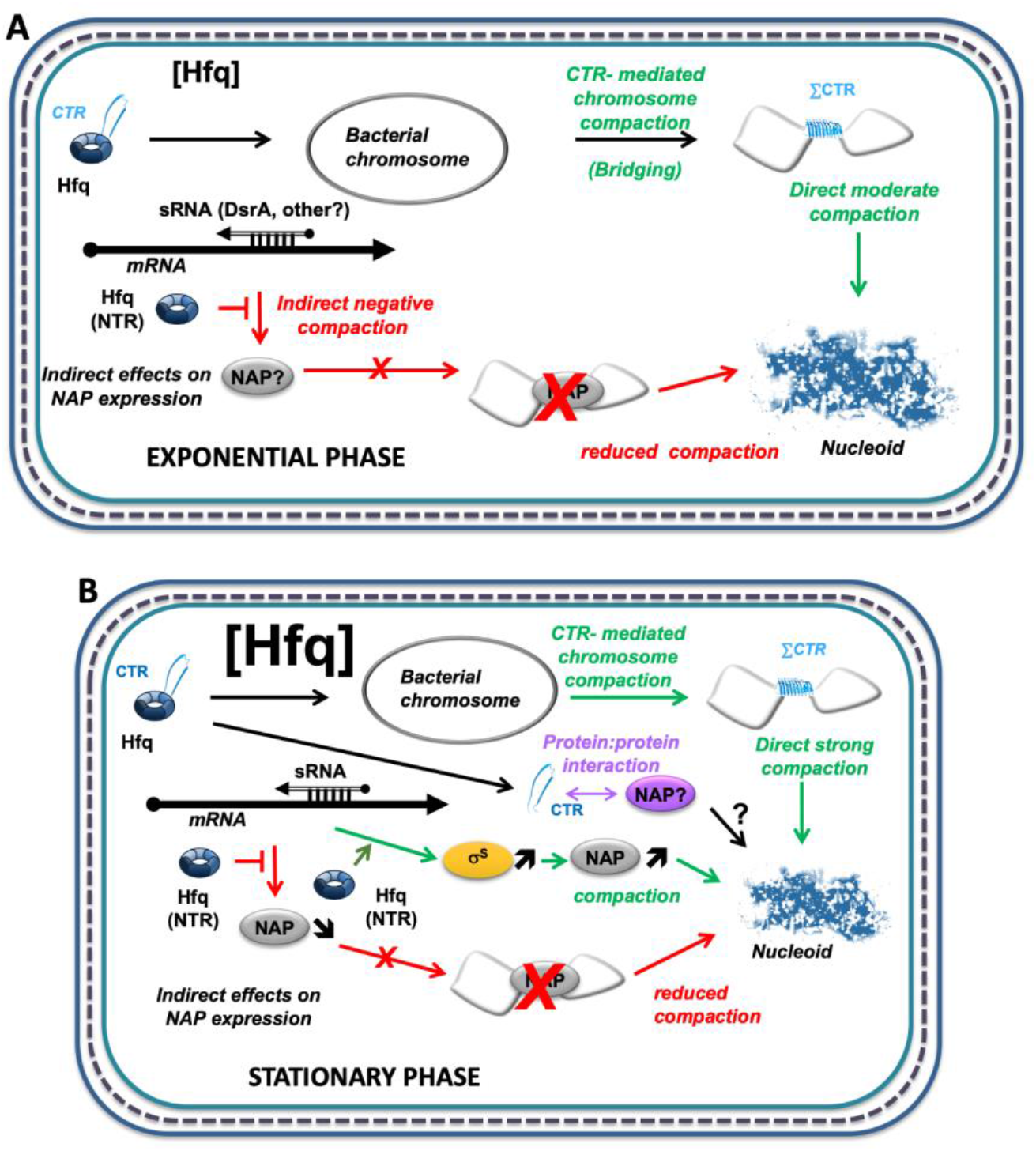
Network of Hfq-dependent regulation controlling DNA compaction. A) Regulation model during the exponential phase. B) Regulation model during the stationary phase. DNA compaction was previously shown to be controlled directly by Hfq-CTR *in vitro*^48^. Briefly, full-length Hfq allows to reduce the nucleoid volume, as expected, but does not significantly change its density (directly related to absorbance). This results from the action of both the NTR and CTR parts of the protein. This effect may be direct *via* the CTR part, or indirect (mainly due to the NTR part). As some NAPs translation such as H-NS are dependent on sRNAs, nucleoid compaction may also be regulated indirectly by Hfq-NTR, most likely negatively^69, 79^. Some NAP may also be under the control of promoters driven by the σ^S^ factor, its expression being itself dependant on sRNAs post-transcriptional positive regulation^67^. sRNA regulators controlling mRNAs are depicted as open arrows; mRNAs as thick black lines; 5’ and 3’ of RNAs are depicted by a “ball and arrow head”, respectively; Hfq-NTR is depicted as a blue toroidal hexamer; Hfq-CTR as a blue amyloid β-strand; σ^S^ as yellow ellipse; NAPs as grey ellipses; chromosomic DNA is represented as a grey line; Nucleoid is represented in blue; positive and negative regulation by Hfq are indicated by arrows and T-shape lines, respectively; dotted line symbolizes Peptidoglycan (PG) between outer (OM) and inner (IM) membranes.

## MATERIAL AND METHODS

All chemicals, unless otherwise stated, were purchased from Sigma-Aldrich.

### Bacteria strains and culture conditions

The bacterial strains used were MG1655 WT (Wild Type, reference strain), MG1655-Δ*hfq* and MG1655-Δ*ctr* (truncated protein with only the first 72 amino-acids)^29^. The *hfq* gene region of these strains have been sequenced to confirm the presence of various *hfq* alleles (*hfq*^+^, Δ*hfq* or Δ*ctr*). For exponential phase cells, bacteria were grown in LB rich media until they reached an OD_600_ of 0.4. For mid-stationary phase, an overnight culture with an OD_600_ 1.5 was analysed. Note that strains do not grow similarly, especially MG1655-Δ*hfq* strain that grows slowly. In addition, the analysis of late stationary phase is difficult in the case of Δ*hfq* strain, due to the effect of Hfq on σ^S^. 1.5 mL of culture were centrifuged, washed in non-supplemented M9 minimal medium and then cells were resuspended in non-supplemented M9 medium and stored at 4°C to stop the growth. Dilutions were performed to reach a final OD_600_ of 0.7 for all strains and conditions of growth. The bacteria were then subsequently processed for cryo-SXT experiments.

### Sample preparation for cryo soft X-ray tomography

Holey carbon-coated gold Quantifoil® Finder grids (Quantifoil Micro Tools, Großlöbichau, Germany) R2/2 (reference: Au G200F1) were glow-discharged using a PELCO EasiGlow™ (Ted Pella, Inc., Redding, CA, USA). Prior deposition on the grids, 10 µL of each sample condition at 0.7 OD_600_ were mixed with 15 µL of 100 nm gold beads (reference: GC100 from BBI solutions) used as fiducial markers for enabling projections alignment to common axis for rotation prior to tomographic reconstruction and 5 µL of M9 media. 5 µL of sample were deposited on the carbon side of the grid, the excess of solution was manually blotted using a Whatman filter paper and then the grid was plunge-frozen into liquid ethane cooled down at - 174°C by liquid nitrogen in a Leica EM CPC cryo-plunger (Leica Microsystems, Wetzlar, Germany). Three grids were prepared per condition, for a total of 18 grids. A few grids were screened with a cryo-transmission electron microscope (JEOL 2200FS) to assess the bacteria concentration and the fiducial distribution on the grid, as well as to verify the absence of ice crystals and evaluate the ice thickness prior to cryo-SXT analysis.

### Cryo soft X-ray tomography

Cryo-SXT dataset were acquired at the ALBA synchrotron Light Source (Barcelona, Spain), on the transmission X-ray microscopy (TXM) beamline (MISTRAL BL09), proposal 2018082926-varluison)^80^.

Samples were mounted in specific holders and transferred to the cryo-TXM experimental vacuum chamber, preserving cryo-conditions throughout the transfer. Once inside the cryo-cooled vacuum chamber the grids were positioned in the focal point and at the axis of rotation of the TXM chamber. Using a visible light microscope on-line with the X-ray microscope areas of interest were selected. Prior to individual bacteria screening, zero-degree soft X-ray projection mosaics were acquired to evaluate sample conditions and to identify best regions on the grids. For our data collection a 25 nm objective Fresnel zone-plate was used to project the transmission signal on a direct illuminated CDD camera (1024 * 1024 pixels, camera pixel size = 13 µm). Acquisitions were performed using X-ray photons in the so-called water-window energy range at 520 eV (corresponding wavelength is 2.38 nm). At this energy, water (Oxygen) is “transparent” (i.e., very low absorption) to X-rays while proteins and DNA (Carbon and Nitrogen) have a strong absorption, thus creating contrast^81^. Tilt-series were collected with an increment of 1° between two projections. The number of projections per tilt-series ranged between 120 and 128 depending on the minimal and maximal tilt-angles that were achieved, which were about ±60°. Magnifications of 1,300X and 1,650X were used, with corresponding image pixel-sizes of 10 nm and 8 nm, respectively. A total of 64 tomographic tilt-series were acquired and distributed as follows: exponential phase WT (12 tomograms), exponential phase Δ*hfq* (14 tomograms), exponential phase Δ*ctr* (10 tomograms), stationary phase WT (9 tomograms), stationary phase Δ*hfq* (9 tomograms) and stationary phase Δ*ctr* (10 tomograms).

### Cryo-SXT tilt-series processing, alignment and reconstruction

To correct the non-uniformity of the X-ray beam, each tilt-series was normalized using an average of 10 images taken without sample, commonly called a flat field (FF). Because the flat field varies over time (it is dependent on the beam intensity of the storage ring), it was measured after each tilt-series collection. The normalisation considered the averaged FF, beam currents and exposure times as denoted in the following equation from^82^:

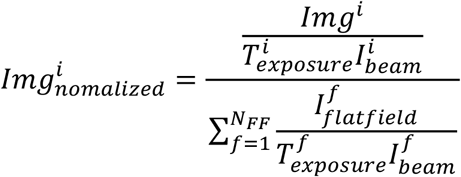

Where *i* is the *i*-th image in the tilt-series, *f* is the *f*-th flat field image and *N*_*FF*_ is the number of FF images used to estimate the average FF, which in our case equal 10 (for each tilt-series). *I* denotes the incident X-ray beam current.

The mass absorption coefficient *μ* is related to the transmitted intensity *I* through a material of density *ρ* and total thickness *d* by:

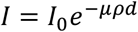

Where *I*_0_ is the intensity impinging on the sample.

The linear absorption coefficient *μ*_*l*_ (*μm*^−1^), which is the reconstructed value in the tomograms, relates to the mass absorption coefficient *μ* (*μm*^2^/*g*) of the material and its density

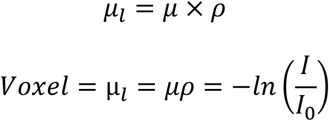

To compute the absorption signal, a negative Napierian logarithm was applied to each normalized tilt-series prior to reconstruction. All tilt-series were processed using a homemade bash script to automate the extraction of the tilt-series from the hdf5 files, absorption tilt-series computation, automated fiducial detection/tracking/alignment using IMOD command lines^83^ and reconstruction of the 3D volumes using SIRT algorithm from Tomo3D software (z = 300 pixels, 20 iterations)^84^.

### Data processing and segmentation

Semi-automatic nucleoid segmentations were performed using a homemade MATLAB (Mathworks Inc., Natick, Massachusetts) and R (https://www.R-project.org/)^85^ combined script. A description of the workflow is present in Supplementary Figure 8. Briefly, the bacteria are cropped in MATLAB. Then, the voxels constituting the nucleoid are found using Gaussian fitting in R. Finally, the segmentation is performed in MATLAB.

### Visualisation of segmented data

The different volumes generated in MATLAB (low-absorbance nucleoid, high-absorbance nucleoid and bacteria volumes) were exported in a format compatible with ChimeraX, which was used for data visualisation^86^. The threshold used for the display of isosurfaces (Figure 4) was the mean value of the voxels (i.e., mean value of membranes and low- and high-absorbance nucleoid voxels). The core domains (Figure 5), corresponding to hollow pocket-like regions inside the nucleoid, are represented using a transparent Gaussian sphere which was designed using the MO 3 command in SPIDER^87^. The size of the sphere was manually adapted to fit the inner diameter of the hollow region.

### Statistical analysis

Because of the low nucleoid volume population sizes and because of their variable sizes (as low as 7 data points for Δ*hfq* exponential phase and stationary phase conditions and up to 19 data points for the WT exponential phase condition), direct population comparison was not relevant. We then used smoothed bootstrap to generate new data points and recover larger populations of comparable (i.e., equal) sizes. Briefly, for each bacteria culture condition, 7 values (i.e., nucleoid volume data points) were randomly picked with replacement. These draws were iteratively repeated 50 times to generate a total of 350 values per bacteria culture condition. We used 7 draws per iteration because it corresponded to the smallest population original size. Indeed, picking more than 7 draws would have more likely generated duplicated draws in the smallest populations (i.e., Δ*hfq* exponential phase and stationary phase conditions) compared to the bigger ones (e.g., the WT exponential phase condition). Moreover, the number of draws had to be consistent. Smoothing was performed by adding *N*(0, σ^2^) random noise to each draw, with 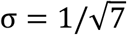.

LAC histograms were computed using R. Histogram counts of each bacterium nucleoid were computed using LAC bins (i.e., intervals) of 0.001. Then for each bin, the counts of each bacterium were converted into the corresponding volume using the voxel size. The bin counts of each bacterium were normalized using 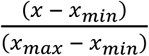 to have a range of volume values in the interval [0, 1]. Finally, the mean and standard deviation of each bin were computed.

For each condition, the mean LAC value of each segmented nucleoid was plotted as a box plot (Figure 2B). To evaluate the statistical significance of the distribution per condition, we first performed Shapiro normality tests on each condition, all of which rejected the non-normal distribution hypothesis. However, because each condition contained less than 30 individuals, Mann-Whitney non-parametric tests were used. A total of nine tests were performed, comparing the two growth phases of each strain (3 tests) and the strains at each growth phase (6 tests). A Bonferroni correction was then applied to compensate for the number of tested hypotheses. We tested *m* = 9 hypotheses with a desired *α* = 0.05 which was corrected to *α*_*Bonferroni*_ = 0.05/9 = 0.00556. The same process was applied to the bootstrapped nucleoid volumes (with this time non-Normal distributions).

### Full length and truncated Hfq relative quantification

Relative quantification of full-length Hfq and CTR-truncated Hfq forms was performed using Western Blot and Dot blot. Briefly, crude extracts at the same OD were diluted serially for both strains. The Δ*hfq* strain was used as control to subtract background. Membranes of blots were successively incubated with anti-Hfq polyclonal antibody from Goat (Origene, Germany), with anti-goat secondary antibody coupled to alkaline phosphatase (Sigma) and revealed with the NBT/BCIP Reagent Kit (Sigma). Intensities of bands were evaluated using ImageJ^88^. Note that the polyclonal anti-Hfq antibodies may recognize slightly less the truncated form of the protein and thus explaining the small differences observed.

## SUPPLEMENTARY DATA

Supplementary Data is available online. Supplementary Figure 1. Tomographic images of exponential phase WT cells. Supplementary Figure 2. Tomographic images of stationary phase WT cells. Supplementary Figure 3. Tomographic images of exponential phase Δ*hfq* cells. Supplementary Figure 4. Tomographic images of stationary phase Δ*hfq* cells. Supplementary Figure 5. Tomographic images of exponential phase Δ*ctr* cells. Supplementary Figure 6. Tomographic images of stationary phase Δ*ctr* cells. Supplementary Figure 7. relative quantification of Hfq in *E. coli* strains. Supplementary Figure 8. Semi-automatic processing of the cryo-SXT data and segmentation of nucleoid voxels.

## ACKNOWLEDGEMENTS

We thank J.M. Carazo and C.O. Sorzano (CNB, Madrid, Spain) for many fruitful discussions. We thank F. Busi (UMR8251, Paris, France) and M. Buckle (ENS, Paris-Saclay, France) for critical reading of the manuscript. We acknowledge ALBA synchrotron (Barcelona, Spain) for provision of synchrotron radiation facilities (proposal 2018082926-varluison) and would like to thank A. Sorrentino for assistance during data collection at the MISTRAL beamline. We acknowledge the Multimodal Imaging Centre at Institut Curie (Orsay, France) for providing access to the cryo-transmission electron microscopy facility.

## FIGURES LEGENDS

**Supplementary Figure 1.**
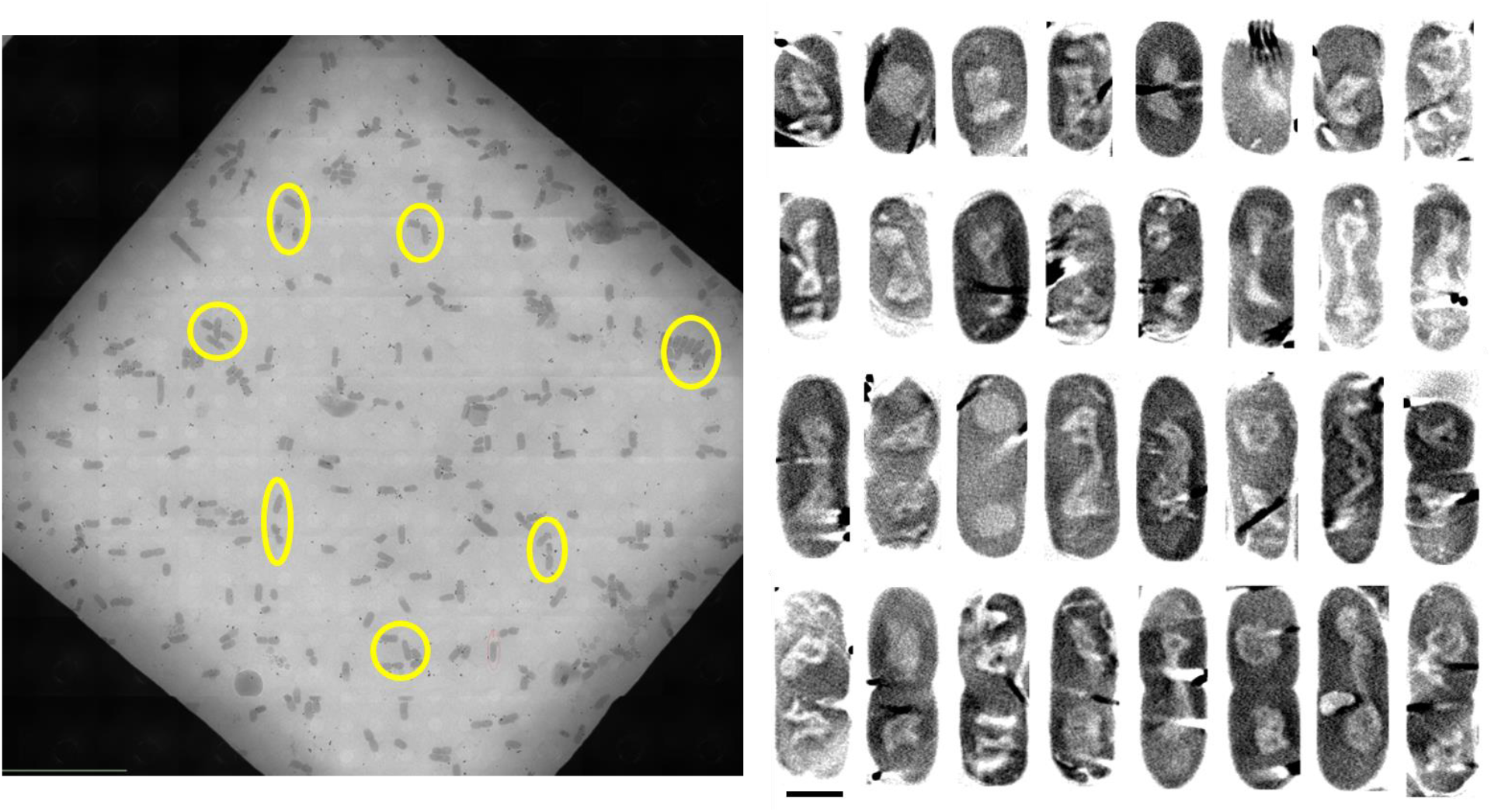
Tomographic reconstructed slices of exponential phase WT cells. Left panel) Low magnification soft-X-ray projection (transmission signal) of a typical grid square showing the distribution of the bacteria on the frozen grid. Yellow circles represent some of the bacteria which were chosen for SXT analysis. Right panel) 10 nm-thick Z-slice of bacteria which were analysed by SXT (images have been extracted from absorbance reconstructed volumes which contrast has been inverted to resemble that of transmission images). Scale bar is 1 µm.

**Supplementary Figure 2.**
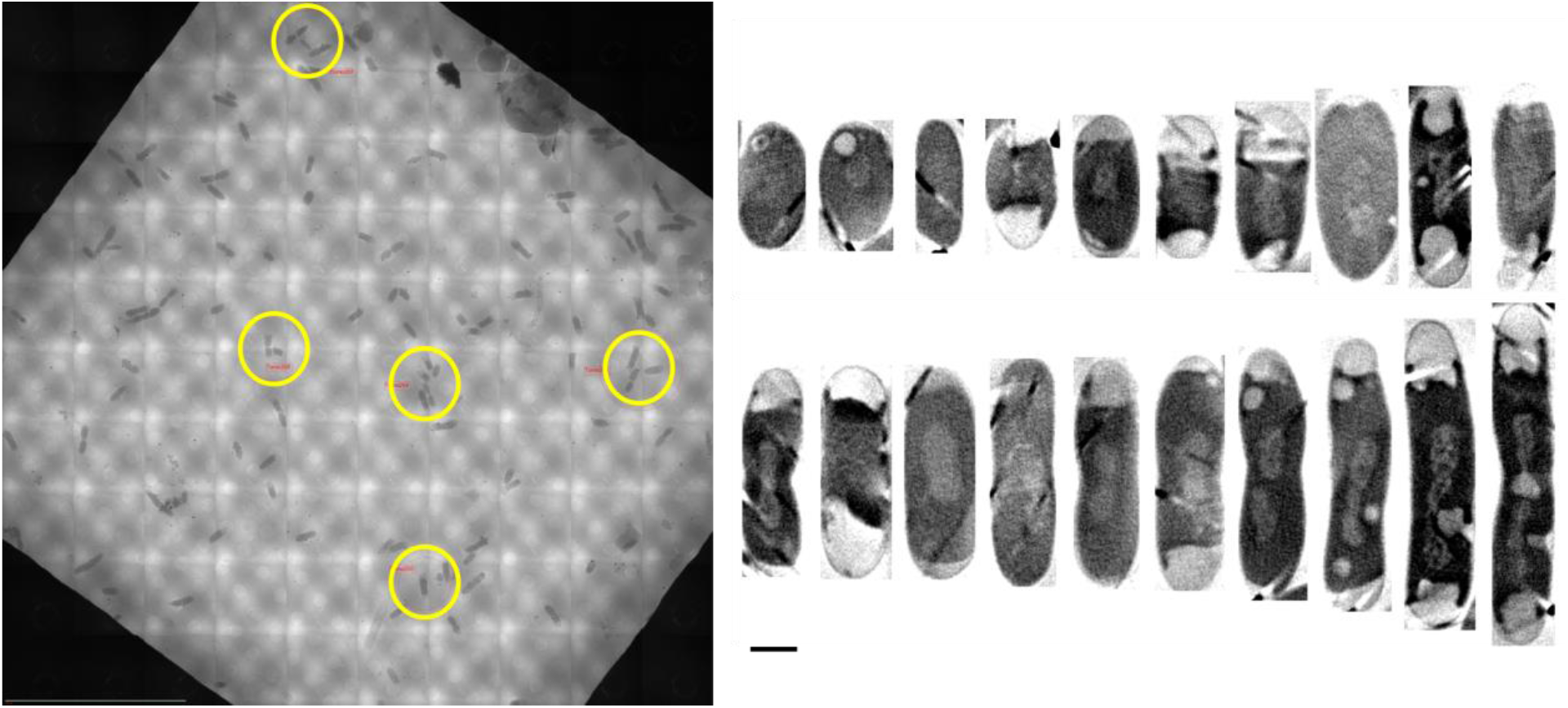
Tomographic reconstruction slices of stationary phase WT cells. Left panel) Low magnification soft-X-ray projection (transmission signal) of a typical grid square showing the distribution of the bacteria on the frozen grid. Yellow circles represent some of the bacteria which were chosen for SXT analysis. Right panel) 10 nm-thick Z-slice of bacteria which were analysed by SXT (images have been extracted from absorbance reconstructed volumes which contrast has been inverted to resemble that of transmission images). Scale bar is 1 µm.

**Supplementary Figure 3.**
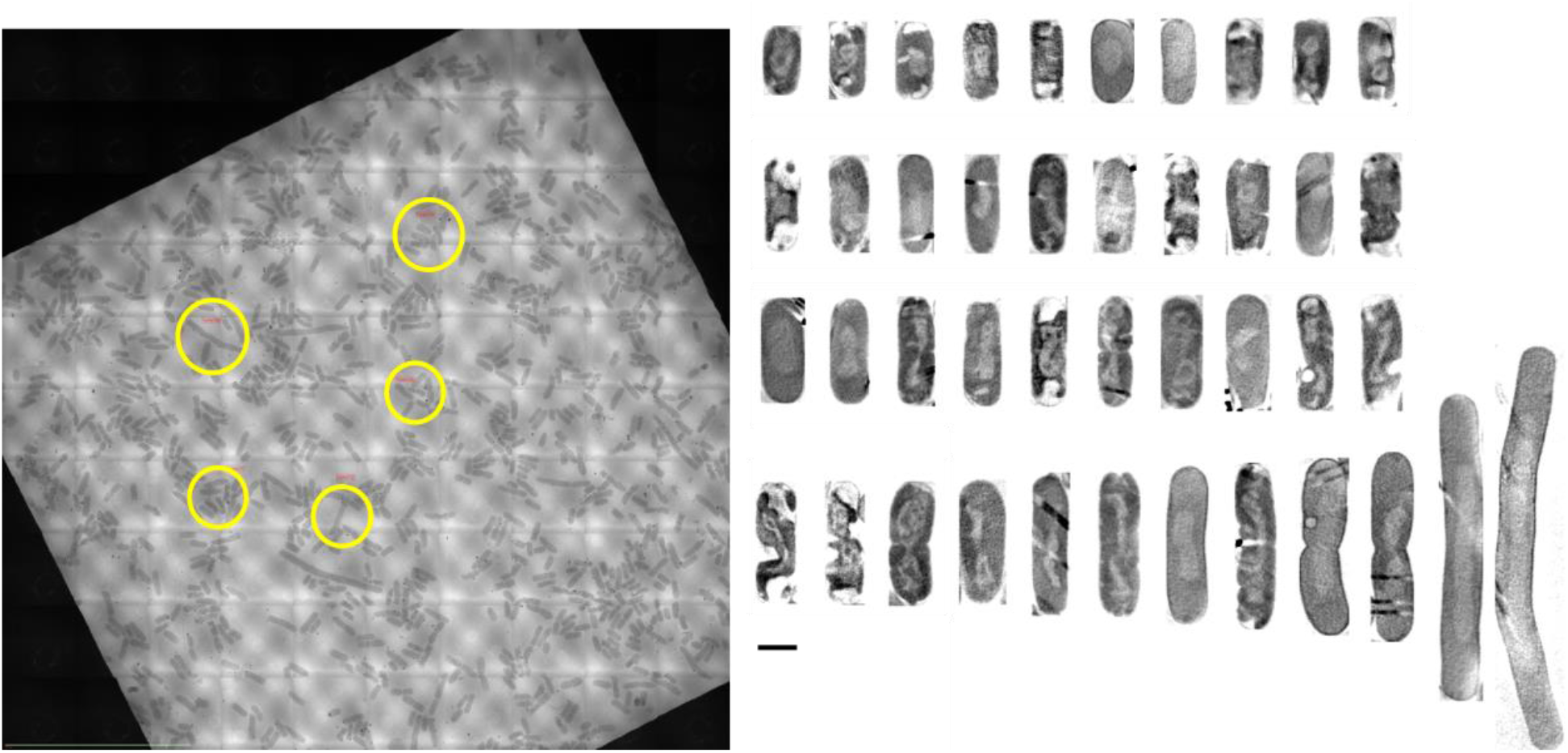
Tomographic reconstruction slices of exponential phase Δ*hfq* cells. Left panel) Low magnification soft-X-ray projection (transmission signal) of a typical grid square showing the distribution of the bacteria on the frozen grid. Yellow circles represent some of the bacteria which were chosen for SXT analysis. Right panel) 10 nm-thick Z-slice of bacteria which were analysed by SXT (images have been extracted from absorbance reconstructed volumes which contrast has been inverted to resemble that of transmission images). Elongated bacteria are visible. Scale bar is 1 µm.

**Supplementary Figure 4.**
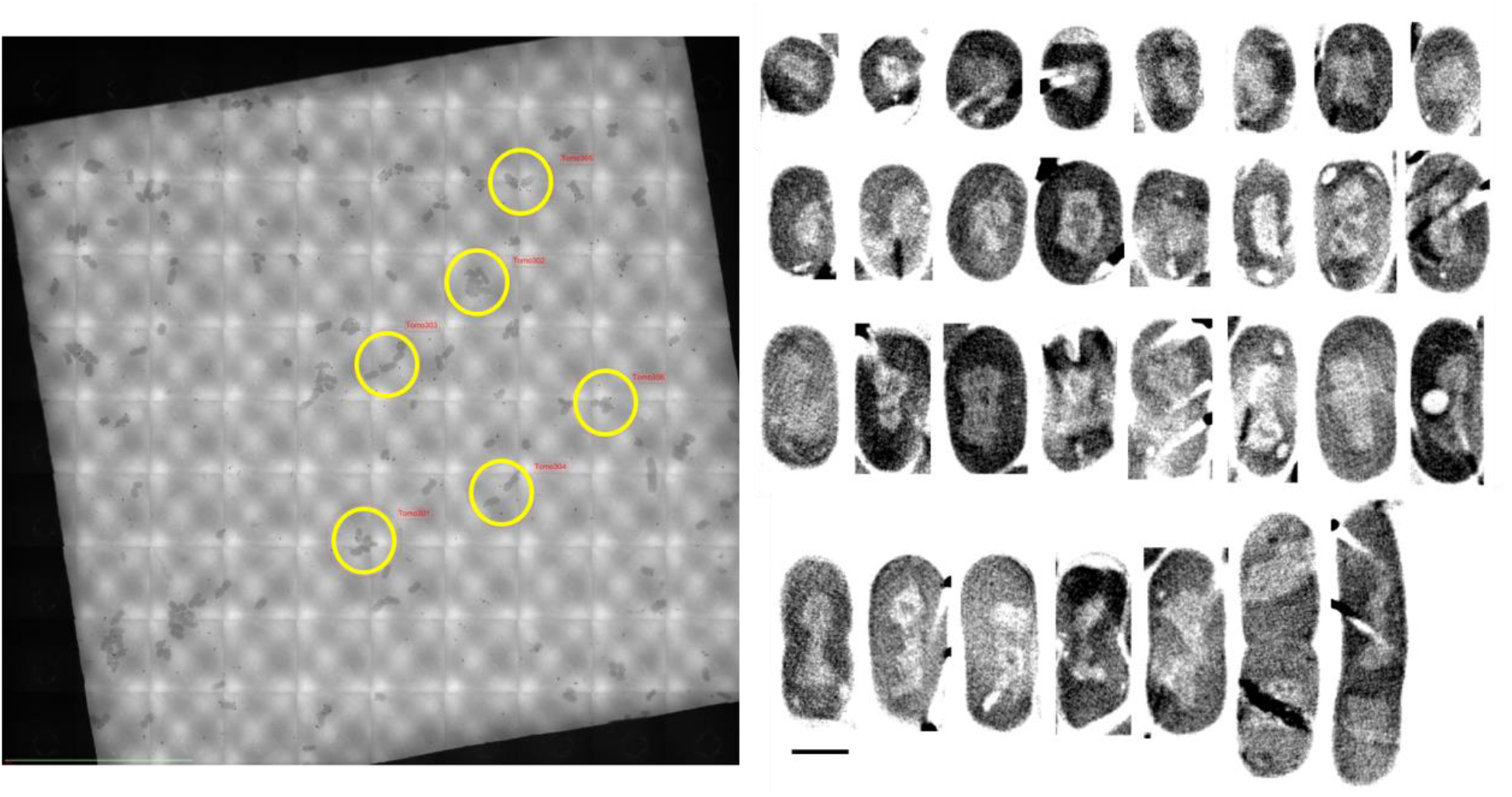
Tomographic reconstruction slices of stationary phase Δ*hfq* cells. Left panel) Low magnification soft-X-ray projection (transmission signal) of a typical grid square showing the distribution of the bacteria on the frozen grid. Yellow circles represent some of the bacteria which were chosen for SXT analysis. Right panel) 10 nm-thick Z-slice of bacteria which were analysed by SXT (images have been extracted from absorbance reconstructed volumes which contrast has been inverted to resemble that of transmission images). Scale bar is 1 µm.

**Supplementary Figure 5.**
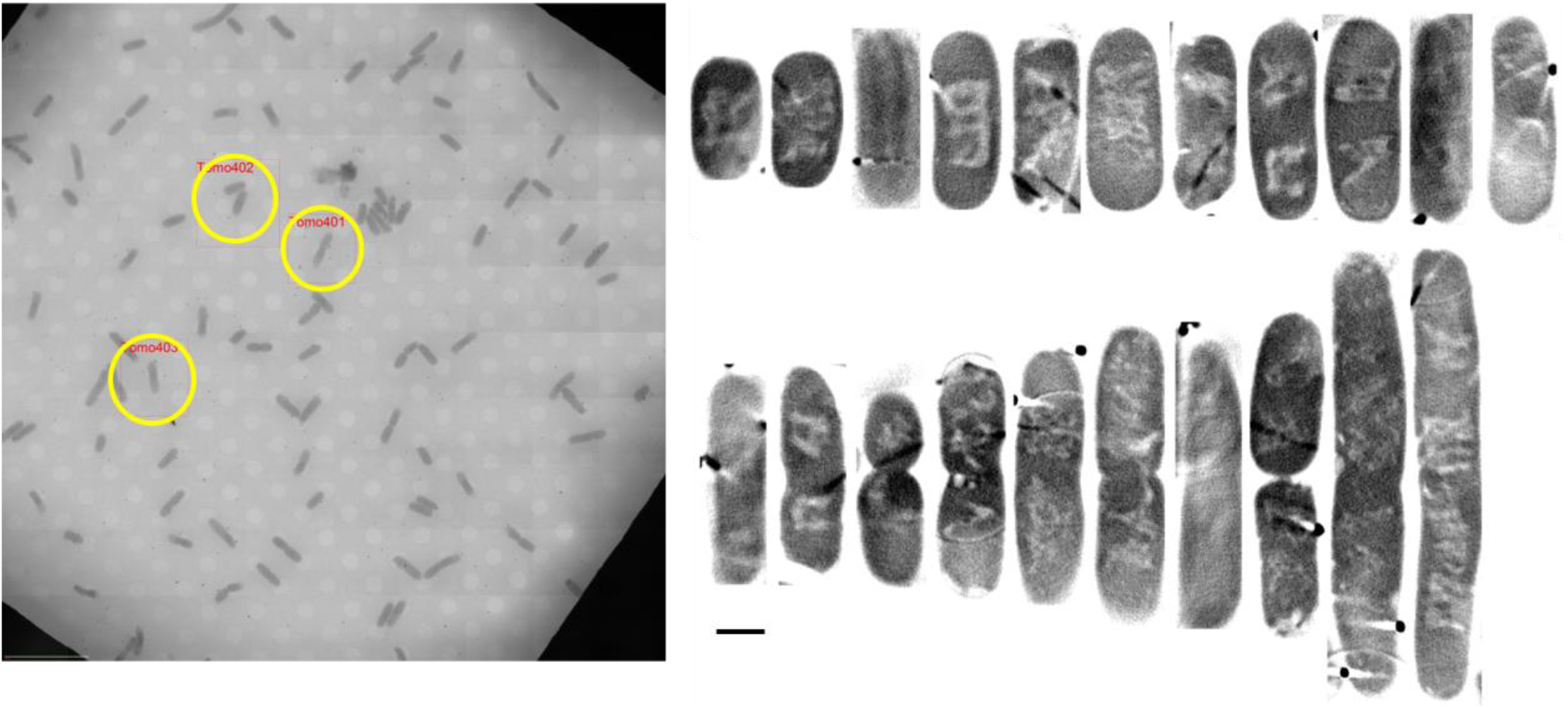
Tomographic reconstruction slices of exponential phase Δ*ctr* cells. Left panel) Low magnification soft-X-ray projection (transmission signal) of a typical grid square showing the distribution of the bacteria on the frozen grid. Yellow circles represent some of the bacteria which were chosen for SXT analysis. Right panel) 10 nm-thick Z-slice of bacteria which were analysed by SXT (images have been extracted from absorbance reconstructed volumes which contrast has been inverted to resemble that of transmission images). Elongated bacteria are visible. Scale bar is 1 µm.

**Supplementary Figure 6.**
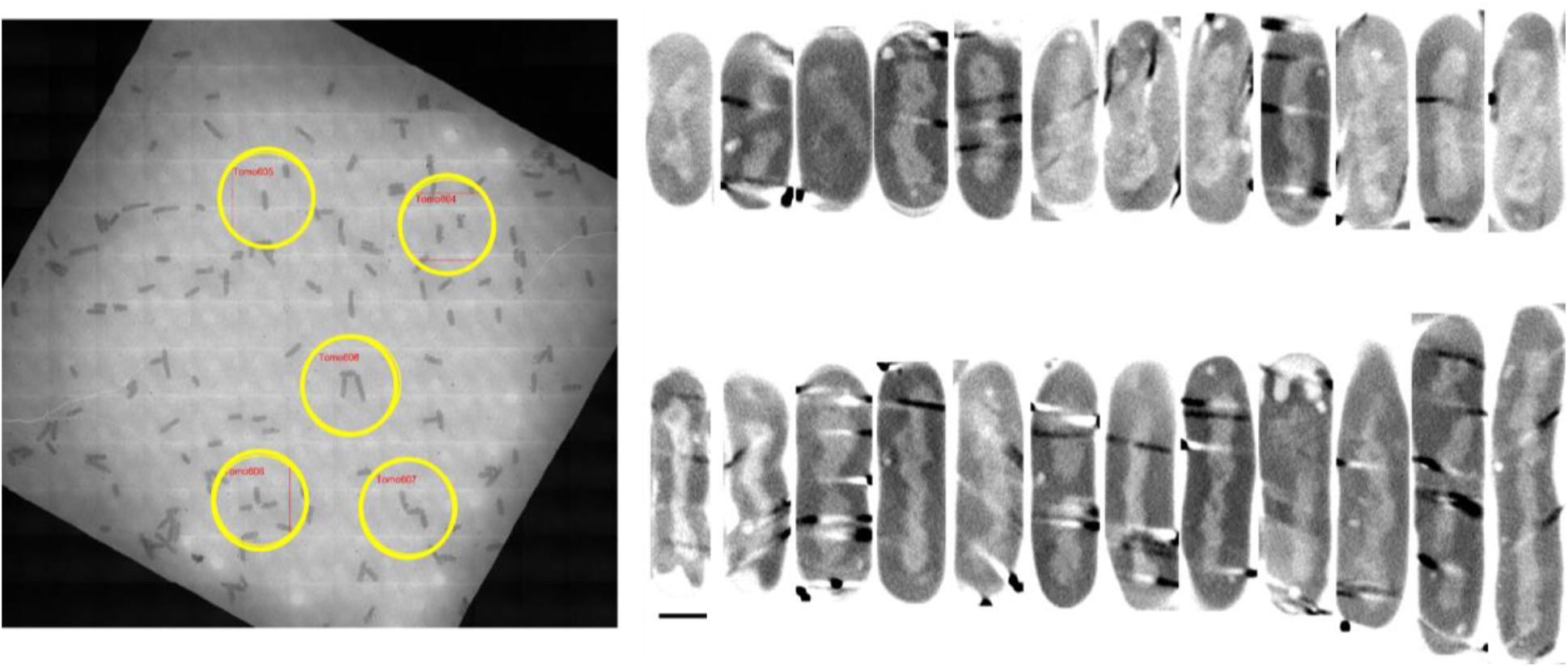
Tomographic reconstruction slices of stationary phase Δ*ctr* cells. Left panel) Low magnification soft-X-ray projection (transmission signal) of a typical grid square showing the distribution of the bacteria on the frozen grid. Yellow circles represent some of the bacteria which were chosen for SXT analysis. Right panel) 10 nm-thick Z-slice of bacteria which were analysed by SXT (images have been extracted from absorbance reconstructed volumes which contrast has been inverted to resemble that of transmission images). Scale bar is 1 µm.

**Supplementary Figure 7.**
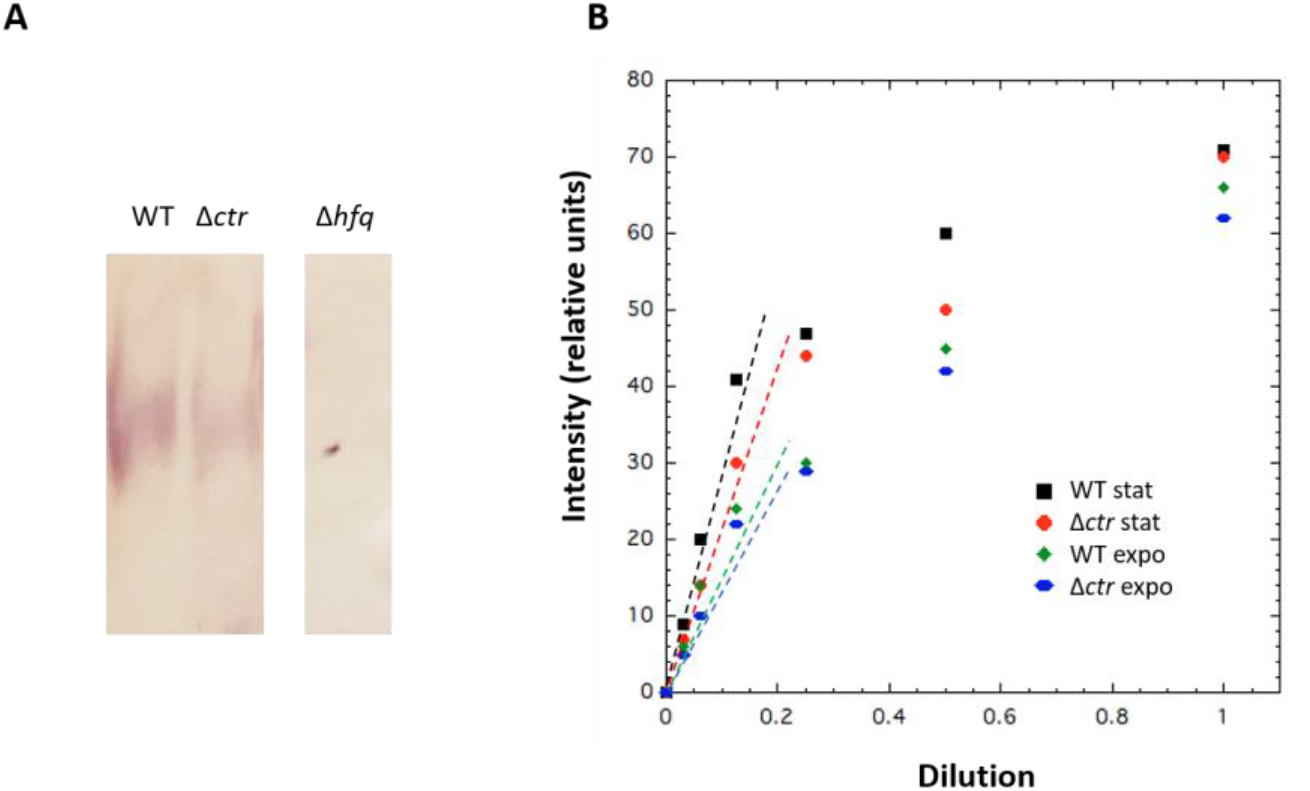
Relative quantification of Hfq in *E. coli* strains. (A) Western Blot (stationary phase); (B) Dot Blot relative quantification. The relative quantification was made in the linear part of the measurements (dotted lines). We confirm that Hfq concentration increases during the stationary phase and that the concentrations of full length (WT) and truncated (Δ*ctr*) forms of Hfq are not significantly different in vivo (polyclonal anti-Hfq antibodies may recognize slightly less the truncated form of the protein and explain the small difference observed).

**Supplementary Figure 8.**
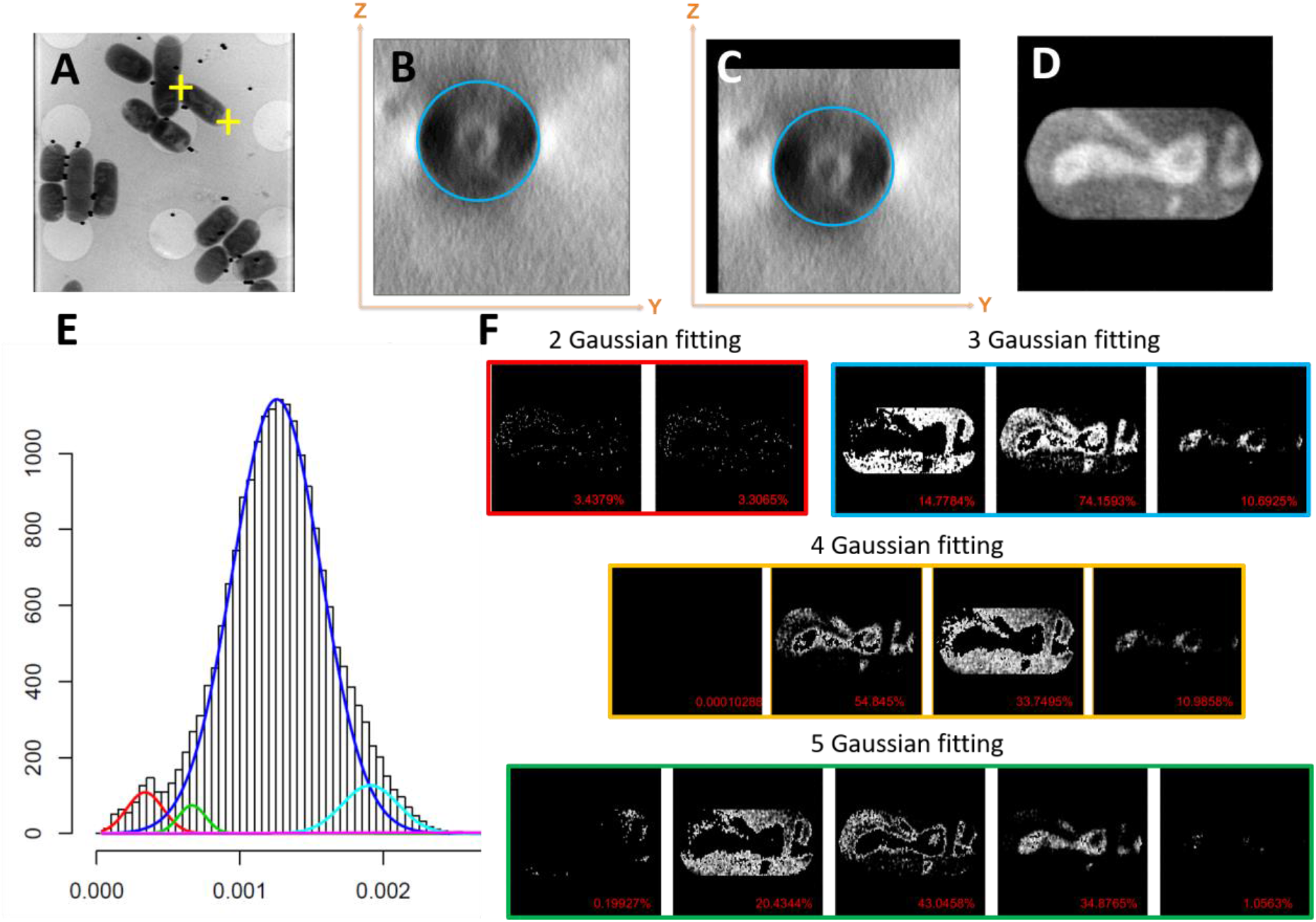
Semi-automatic processing of the cryo-SXT data and segmentation of nucleoid voxels. The processing of the cryo-SXT data to perform the segmentation of the nucleoid was performed in MATLAB and R. This supplementary figure presents the whole workflow. A) In MATLAB, the bacterium poles are clicked on the central Z-section of each reconstructed tomogram, generating a pair of XY coordinates (yellow crosses). A sub-tomogram containing the full bacterium is then cropped from the original 3D volume and rotated to place its long axis (i.e., the bacterium length) horizontally. B) Following that, an average of 5 central slices (along the X-axis) is displayed permitting to select a circle (or an ellipse) fitting the bacterium transversal slice. C) These two steps permit to compute the length, centre and width of the bacterium, enabling the centring of the bacterium inside the 3D volume. D) Using these metrics, the script fits a sphero-cylinder on the bacterium and each voxel located outside the sphero-cylinder is set to NaN so that these voxels can be discarded during the next computing steps. Then a 3D median filter (kernel [3, 3, 3]) is applied and the voxel values are exported. The purpose of the median filter is to smooth the voxel value distribution for further analysis. This procedure allowed to extract 30 exponential phase WT bacteria, 66 exponential phase Δ*hfq* bacteria, 18 exponential phase Δ*ctr* bacteria, 17 stationary phase WT bacteria, 27 stationary phase Δ*hfq* bacteria and 24 stationary phase Δ*ctr* bacteria. The next step is performed in R. Previously extracted voxel values are imported in R and analysed using the Mixtools package^89^. E) This package allows to detect multimodal distributions by Gaussian fitting of several curves (in the present study we fitted up to 5 Gaussians). F) Each fitted Gaussian curve represents a sub-population of voxels with similar grey values, which in some cases, corresponds to a particular compartment of the bacterial cell (e.g. membranes, cytoplasm, nucleoid or gold beads). Using this automatic selection of bacterial compartments, we were able to accurately determine which pixels constituted the nucleoid. When this analysis did not generate a clear-cut dissection of the bacterial compartments, the corresponding bacterium was discarded. Clear-cut dissection was generally not achieved when *i*) focus changed during data collection, *ii*) alignment was poor by lack of fiducials or *iii*) too many gold beads where present next to the bacterium, drastically compressing the contrast range. After this rigorous selection, 19 exponential phase WT bacteria, 7 exponential phase Δ*hfq* bacteria, 10 exponential phase Δ*ctr* bacteria, 11 stationary phase WT bacteria, 7 stationary phase Δ*hfq* bacteria and 12 stationary phase Δ*ctr* bacteria were automatically successfully segmented. After determining which voxels constituted the nucleoid, it was then possible to compute the volume occupied by the nucleoid for each bacterium. To highlight the variability of the LAC values inside each nucleoid, we chose to perform iterative grayscale erosion to equally split the low-absorbance nucleoid from the high-absorbance one in MATLAB. After erosion, the low-absorbance nucleoid and the high-absorbance one occupied the same volume (i.e., each of them occupied half the volume of the whole nucleoid). Two volumes were then generated, one corresponding to the low-absorbance nucleoid, the other corresponding to the high-absorbance one.

## REFERENCES

1. Sun, X.; Zhulin, I.; Wartell, R. M., Predicted structure and phyletic distribution of the RNA-binding protein Hfq. Nucleic Acids Res 2002, 30 (17), 3662–71.

2. Bouloc, P.; Repoila, F., Fresh layers of RNA-mediated regulation in Gram-positive bacteria. Curr Opin Microbiol 2016, 30, 30–35.

3. Franze de Fernandez, M.T.; Eoyang, L.; August, J. T., Factor fraction required for the synthesis of bacteriophage Qbeta-RNA. Nature 1968, 219 (5154), 588–90.

4. Tsui, H. C.; Leung, H. C.; Winkler, M. E., Characterization of broadly pleiotropic phenotypes caused by an hfq insertion mutation in Escherichia coli K-12. Mol. Microbiol. 1994, 13, 35–49.

5. Rajkowitsch, L.; Chen, D.; Stampfl, S.; Semrad, K.; Waldsich, C.; Mayer, O.; Jantsch, M. F.; Konrat, R.; Blasi, U.; Schroeder, R., RNA chaperones, RNA annealers and RNA helicases. RNA Biol 2007, 4 (3), 118–30.

6. Arluison, V.; Hohng, S.; Roy, R.; Pellegrini, O.; Regnier, P.; Ha, T., Spectroscopic observation of RNA chaperone activities of Hfq in post-transcriptional regulation by a small non-coding RNA. Nucleic Acids Res 2007, 35 (3), 999–1006.

7. Vogel, J.; Luisi, B. F., Hfq and its constellation of RNA. Nat Rev Microbiol 2011, 9 (8), 578–89.

8. Aiba, H., Mechanism of RNA silencing by Hfq-binding small RNAs. Curr Opin Microbiol 2007, 10 (2), 134–9.

9. McCullen, C. A.; Benhammou, J. N.; Majdalani, N.; Gottesman, S., Mechanism of positive regulation by DsrA and RprA small noncoding RNAs: pairing increases translation and protects rpoS mRNA from degradation. J Bacteriol 2010, 192 (21), 5559–71.

10. Udekwu, K. I.; Darfeuille, F.; Vogel, J.; Reimegard, J.; Holmqvist, E.; Wagner, E. G., Hfq-dependent regulation of OmpA synthesis is mediated by an antisense RNA. Genes Dev 2005, 19 (19), 2355–66.

11. Gottesman, S.; McCullen, C. A.; Guillier, M.; Vanderpool, C. K.; Majdalani, N.; Benhammou, J.; Thompson, K. M.; FitzGerald, P. C.; Sowa, N. A.; FitzGerald, D. J., Small RNA regulators and the bacterial response to stress. Cold Spring Harb Symp Quant Biol 2006, 71, 1–11.

12. Gottesman, S., Trouble is coming: Signaling pathways that regulate general stress responses in bacteria. The Journal of biological chemistry 2019, 294 (31), 11685–11700.

13. Kendall, M. M.; Gruber, C. C.; Rasko, D. A.; Hughes, D. T.; Sperandio, V., Hfq virulence regulation in enterohemorrhagic Escherichia coli O157:H7 strain 86-24. J Bacteriol 2011, 193 (24), 6843–51.

14. Porcheron, G.; Dozois, C. M., Interplay between iron homeostasis and virulence: Fur and RyhB as major regulators of bacterial pathogenicity. Vet Microbiol 2015, 179 (1-2), 2–14.

15. Ding, Y.; Davis, B. M.; Waldor, M. K., Hfq is essential for Vibrio cholerae virulence and downregulates sigma expression. Mol Microbiol 2004, 53 (1), 345–54.

16. Gaffke, L.; Kubiak, K.; Cyske, Z.; Wegrzyn, G., Differential Chromosome- and Plasmid-Borne Resistance of Escherichia coli hfq Mutants to High Concentrations of Various Antibiotics. Int J Mol Sci 2021, 22 (16).

17. Lenz, D. H.; Mok, K. C.; Lilley, B. N.; Kulkarni, R. V.; Wingreen, N. S.; Bassler, B. L., The small RNA chaperone Hfq and multiple small RNAs control quorum sensing in Vibrio harveyi and Vibrio cholerae. Cell 2004, 118 (1), 69–82.

18. Wilusz, C. J.; Wilusz, J., Eukaryotic Lsm proteins: lessons from bacteria. Nat Struct Mol Biol 2005, 12 (12), 1031–6.

19. Brennan, R. G.; Link, T. M., Hfq structure, function and ligand binding. Curr Opin Microbiol 2007, 10 (2), 125–33.

20. Robinson, K. E.; Orans, J.; Kovach, A. R.; Link, T. M.; Brennan, R. G., Mapping Hfq-RNA interaction surfaces using tryptophan fluorescence quenching. Nucleic Acids Res 2014, 42 (4), 2736–49.

21. Sauer, E.; Schmidt, S.; Weichenrieder, O., Small RNA binding to the lateral surface of Hfq hexamers and structural rearrangements upon mRNA target recognition. Proc Natl Acad Sci U S A 2012, 109 (24), 9396–401.

22. Updegrove, T. B.; Zhang, A.; Storz, G., Hfq: the flexible RNA matchmaker. Curr Opin Microbiol 2016, 30, 133–8.

23. Arluison, V.; Folichon, M.; Marco, S.; Derreumaux, P.; Pellegrini, O.; Seguin, J.; Hajnsdorf, E.; Regnier, P., The C-terminal domain of Escherichia coli Hfq increases the stability of the hexamer. Eur J Biochem 2004, 271 (7), 1258–65.

24. Wen, B.; Wang, W.; Zhang, J.; Gong, Q.; Shi, Y.; Wu, J.; Zhang, Z., Structural and dynamic properties of the C-terminal region of the Escherichia coli RNA chaperone Hfq: integrative experimental and computational studies. Phys Chem Chem Phys 2017, 19 (31), 21152–21164.

25. Orans, J.; Kovach, A. R.; Hoff, K. E.; Horstmann, N. M.; Brennan, R. G., Crystal structure of an Escherichia coli Hfq Core (residues 2-69)-DNA complex reveals multifunctional nucleic acid binding sites. Nucleic acids res 2020, 48 (7), 3987–3997.

26. Dimastrogiovanni, D.; Frohlich, K. S.; Bandyra, K. J.; Bruce, H. A.; Hohensee, S.; Vogel, J.; Luisi, B. F., Recognition of the small regulatory RNA RydC by the bacterial Hfq protein. Elife 2014, 3.

27. Fortas, E.; Piccirilli, F.; Malabirade, A.; Militello, V.; Trepout, S.; Marco, S.; Taghbalout, A.; Arluison, V., New insight into the structure and function of Hfq C-terminus. Biosci Rep 2015, 35 (2).

28. Partouche, D.; Militello, V.; Gomez-Zavaglia, A.; Wien, F.; Sandt, C.; Arluison, V., In Situ Characterization of Hfq Bacterial Amyloid: A Fourier-Transform Infrared Spectroscopy Study. Pathogens 2019, 8 (1).

29. Malabirade, A.; Partouche, D.; El Hamoui, O.; Turbant, F.; Geinguenaud, F.; Recouvreux, P.; Bizien, T.; Busi, F.; Wien, F.; Arluison, V., Revised role for Hfq bacterial regulator on DNA topology. Sci Rep 2018, 8 (1), 16792.

30. Olsen, A. S.; Moller-Jensen, J.; Brennan, R. G.; Valentin-Hansen, P., C-Terminally truncated derivatives of Escherichia coli Hfq are proficient in riboregulation. J Mol Biol 2010, 404 (2), 173–82.

31. Vecerek, B.; Rajkowitsch, L.; Sonnleitner, E.; Schroeder, R.; Blasi, U., The C-terminal domain of Escherichia coli Hfq is required for regulation. Nucleic Acids Res 2008, 36 (1), 133–43.

32. Vincent, H. A.; Henderson, C. A.; Stone, C. M.; Cary, P. D.; Gowers, D. M.; Sobott, F.; Taylor, J. E.; Callaghan, A. J., The low-resolution solution structure of Vibrio cholerae Hfq in complex with Qrr1 sRNA. Nucleic Acids Res 2012, 40 (17), 8698–710.

33. Salim, N. N.; Faner, M. A.; Philip, J. A.; Feig, A. L., Requirement of upstream Hfq-binding (ARN)x elements in glmS and the Hfq C-terminal region for GlmS upregulation by sRNAs GlmZ and GlmY. Nucleic Acids Res 2012, 40 (16), 8021–32.

34. Turbant, F.; Wu, P.; Wien, F.; Arluison, V., The Amyloid Region of Hfq Riboregulator Promotes DsrA:rpoS RNAs Annealing. Biology (Basel) 2021, 10 (9).

35. Takada, A.; Wachi, M.; Kaidow, A.; Takamura, M.; Nagai, K., DNA binding properties of the hfq gene product of Escherichia coli. Biochem. Biophys. Res. Com. 1997, 236, 576–579.

36. Dorman, C. J., Function of nucleoid-associated proteins in chromosome structuring and transcriptional regulation. J Mol Microbiol Biotechnol 2014, 24 (5-6), 316–31.

37. Azam, T. A.; Ishihama, A., Twelve species of the nucleoid-associated protein from Escherichia coli. Sequence recognition specificity and DNA binding affinity. J Biol Chem 1999, 274 (46), 33105–13.

38. Dillon, S. C.; Dorman, C. J., Bacterial nucleoid-associated proteins, nucleoid structure and gene expression. Nature reviews Microbiology 2010, 8 (3), 185–95.

39. Dame, R. T.; Rashid, F.-Z. M.; Grainger, D. C., Chromosome organization in bacteria: mechanistic insights into genome structure and function. Nature reviews Genetics 2020, 21 (4), 227–242.

40. Diestra, E.; Cayrol, B.; Arluison, V.; Risco, C., Cellular electron microscopy imaging reveals the localization of the Hfq protein close to the bacterial membrane. PLoS One 2009, 4 (12), e8301.

41. Azam, T. A.; Hiraga, S.; Ishihama, A., Two types of localization of the DNA-binding proteins within the Escherichia coli nucleoid. Genes Cells 2000, 5 (8), 613–26.

42. Milles, S.; Huy Bui, K.; Koehler, C.; Eltsov, M.; Beck, M.; Lemke, E. A., Facilitated aggregation of FG nucleoporins under molecular crowding conditions. EMBO Rep 2013, 14 (2), 178–83.

43. Wiggins, P. A.; Dame, R. T.; Noom, M. C.; Wuite, G. J., Protein-mediated molecular bridging: a key mechanism in biopolymer organization. Biophys J 2009, 97 (7), 1997–2003.

44. Geinguenaud, F.; Calandrini, V.; Teixeira, J.; Mayer, C.; Liquier, J.; Lavelle, C.; Arluison, V., Conformational transition of DNA bound to Hfq probed by infrared spectroscopy. Phys Chem Chem Phys 2011, 13 (3), 1222–9.

45. Jiang, K.; Zhang, C.; Guttula, D.; Liu, F.; van Kan, J. A.; Lavelle, C.; Kubiak, K.; Malabirade, A.; Lapp, A.; Arluison, V.; van der Maarel, J. R., Effects of Hfq on the conformation and compaction of DNA. Nucleic Acids Res 2015, 43 (8), 4332–41.

46. El Hamoui, O.; Yadav, I.; Radiom, M.; Wien, F.; Berret, J.-F.; van der Maarel, J. R. C.; Arluison, V., Interactions between DNA and the Hfq Amyloid-like Region Trigger a Viscoelastic Response. Biomacromolecules 2020, 21 (9), 3668–3677.

47. Skoko, D.; Yoo, D.; Bai, H.; Schnurr, B.; Yan, J.; McLeod, S. M.; Marko, J. F.; Johnson, R. C., Mechanism of chromosome compaction and looping by the Escherichia coli nucleoid protein Fis. Journal of molecular biology 2006, 364 (4), 777–98.

48. Malabirade, A.; Jiang, K.; Kubiak, K.; Diaz-Mendoza, A.; Liu, F.; van Kan, J. A.; Berret, J. F.; Arluison, V.; van der Maarel, J. R. C., Compaction and condensation of DNA mediated by the C-terminal domain of Hfq. Nucleic Acids Res 2017, 45 (12), 7299–7308.

49. Updegrove, T. B.; Correia, J. J.; Galletto, R.; Bujalowski, W.; Wartell, R. M., E. coli DNA associated with isolated Hfq interacts with Hfq’s distal surface and C-terminal domain. Biochim Biophys Acta 2010, 1799 (8), 588–96.

50. Tolstorukov, M. Y.; Virnik, K. M.; Adhya, S.; Zhurkin, V. B., A-tract clusters may facilitate DNA packaging in bacterial nucleoid. Nucleic Acids Res 2005, 33 (12), 3907–18.

51. Aiyar, S. E.; Gourse, R. L.; Ross, W., Upstream A-tracts increase bacterial promoter activity through interactions with the RNA polymerase alpha subunit. Proc Natl Acad Sci U S A 1998, 95 (25), 14652–7.

52. Cech, G. M.; Pakula, B.; Kamrowska, D.; Wegrzyn, G.; Arluison, V.; Szalewska-Palasz, A., Hfq protein deficiency in Escherichia coli affects ColE1-like but not lambda plasmid DNA replication. Plasmid 2014, 73, 10–5.

53. Haniford, D. B.; Ellis, M. J., Transposons Tn10 and Tn5. Microbiol Spectr 2015, 3 (1), MDNA3-0002-2014.

54. Le Derout, J.; Boni, I. V.; Regnier, P.; Hajnsdorf, E., Hfq affects mRNA levels independently of degradation. BMC Mol Biol 2010, 11, 17.

55. Parekh, V. J.; Niccum, B. A.; Shah, R.; Rivera, M. A.; Novak, M. J.; Geinguenaud, F.; Wien, F.; Arluison, V.; Sinden, R. R., Role of Hfq in Genome Evolution: Instability of G-Quadruplex Sequences in E. coli. Microorganisms 2019, 8 (1).

56. Chen, J.; Gottesman, S., Hfq links translation repression to stress-induced mutagenesis in E. coli. Genes & development 2017, 31 (13), 1382–1395.

57. Oton, J.; Pereiro, E.; Perez-Berna, A. J.; Millach, L.; Sorzano, C. O.; Marabini, R.; Carazo, J. M., Characterization of transfer function, resolution and depth of field of a soft X-ray microscope applied to tomography enhancement by Wiener deconvolution. Biomed Opt Express 2016, 7 (12), 5092–5103.

58. Hoff, K. A., Rapid and simple method for double staining of bacteria with 4’,6-diamidino-2-phenylindole and fluorescein isothiocyanate-labeled antibodies. Appl Environ Microbiol 1988, 54 (12), 2949–52.

59. Wang, W.; Li, G. W.; Chen, C.; Xie, X. S.; Zhuang, X., Chromosome organization by a nucleoid-associated protein in live bacteria. Science 2011, 333 (6048), 1445–9.

60. Bettridge, K.; Verma, S.; Weng, X.; Adhya, S.; Xiao, J., Single-molecule tracking reveals that the nucleoid-associated protein HU plays a dual role in maintaining proper nucleoid volume through differential interactions with chromosomal DNA. Mol Microbiol 2021, 115 (1), 12–27.

61. Hammel, M.; Amlanjyoti, D.; Reyes, F. E.; Chen, J. H.; Parpana, R.; Tang, H. Y.; Larabell, C. A.; Tainer, J. A.; Adhya, S., HU multimerization shift controls nucleoid compaction. Sci Adv 2016, 2 (7), e1600650.

62. Groen, J.; Conesa, J. J.; Valcarcel, R.; Pereiro, E., The cellular landscape by cryo soft X-ray tomography. Biophys Rev 2019, 611–619.

63. Mandala, V. S.; Loh, D. M.; Shepard, S. M.; Geeson, M. B.; Sergeyev, I. V.; Nocera, D. G.; Cummins, C. C.; Hong, M., Bacterial Phosphate Granules Contain Cyclic Polyphosphates: Evidence from (31)P Solid-State NMR. J Am Chem Soc 2020, 142 (43), 18407–18421.

64. Racki, L. R.; Tocheva, E. I.; Dieterle, M. G.; Sullivan, M. C.; Jensen, G. J.; Newman, D. K., Polyphosphate granule biogenesis is temporally and functionally tied to cell cycle exit during starvation in Pseudomonas aeruginosa. Proc Natl Acad Sci U S A 2017, 114 (12), E2440–E2449.

65. Pallerla, S. R.; Knebel, S.; Polen, T.; Klauth, P.; Hollender, J.; Wendisch, V. F.; Schoberth, S. M., Formation of volutin granules in Corynebacterium glutamicum. FEMS Microbiol Lett 2005, 243 (1), 133–40.

66. Zhu, Y.; Mustafi, M.; Weisshaar, J. C., Biophysical Properties of Escherichia coli Cytoplasm in Stationary Phase by Superresolution Fluorescence Microscopy. mBio 2020, 11 (3).

67. Battesti, A.; Majdalani, N.; Gottesman, S., The RpoS-Mediated General Stress Response in Escherichia coli (*). Annu Rev Microbiol 2011, 65, 189–213.

68. Lee, S. Y.; Lim, C. J.; Droge, P.; Yan, J., Regulation of Bacterial DNA Packaging in Early Stationary Phase by Competitive DNA Binding of Dps and IHF. Sci Rep 2015, 5, 18146.

69. Majdalani, N.; Cunning, C.; Sledjeski, D.; Elliott, T.; Gottesman, S., DsrA RNA regulates translation of RpoS message by an anti-antisense mechanism, independent of its action as an antisilencer of transcription. Proc Natl Acad Sci U S A 1998, 95 (21), 12462–7.

70. Lease, R. A.; Belfort, M., Riboregulation by DsrA RNA: trans-actions for global economy. Mol Microbiol 2000, 38 (4), 667–72.

71. Lalaouna, D.; Masse, E., The spectrum of activity of the small RNA DsrA: not so narrow after all. Curr Genet 2016, 62 (2), 261–4.

72. Gao, Y.; Foo, Y. H.; Winardhi, R. S.; Tang, Q.; Yan, J.; Kenney, L. J., Charged residues in the H-NS linker drive DNA binding and gene silencing in single cells. Proc Natl Acad Sci U S A 2017, 114 (47), 12560–12565.

73. Khosraviani, N.; Ostrowski, L. A.; Mekhail, K., Roles for Non-coding RNAs in Spatial Genome Organization. Front Cell Dev Biol 2019, 7, 336.

74. Kajitani, M.; Ishihama, A., Identification and sequence determination of the host factor gene for bacteriophage Qb. Nucl. Acids Res. 1991, 9, 1063–1066.

75. Remesh, S. G.; Verma, S. C.; Chen, J. H.; Ekman, A. A.; Larabell, C. A.; Adhya, S.; Hammel, M., Nucleoid remodeling during environmental adaptation is regulated by HU-dependent DNA bundling. Nat Commun 2020, 11 (1), 2905.

76. Valens, M.; Penaud, S.; Rossignol, M.; Cornet, F.; Boccard, F., Macrodomain organization of the Escherichia coli chromosome. EMBO J 2004, 23 (21), 4330–41.

77. Espeli, O.; Boccard, F., Organization of the Escherichia coli chromosome into macrodomains and its possible functional implications. J Struct Biol 2006, 156 (2), 304–10.

78. Holowka, J.; Zakrzewska-Czerwinska, J., Nucleoid Associated Proteins: The Small Organizers That Help to Cope With Stress. Front Microbiol 2020, 11, 590.

79. Lalaouna, D.; Morissette, A.; Carrier, M. C.; Masse, E., DsrA regulatory RNA represses both hns and rbsD mRNAs through distinct mechanisms in Escherichia coli. Mol Microbiol 2015, 98 (2), 357–69.

80. Sorrentino, A.; Nicolas, J.; Valcarcel, R.; Chichon, F. J.; Rosanes, M.; Avila, J.; Tkachuk, A.; Irwin, J.; Ferrer, S.; Pereiro, E., MISTRAL: a transmission soft X-ray microscopy beamline for cryo nano-tomography of biological samples and magnetic domains imaging. J Synchrotron Radiat 2015, 22 (4), 1112–7.

81. Wolter, H., Spiegelsysteme streifenden Einfalls als abbildende Optiken für Röntgenstrahlen. Wiley: 1952; Vol. 445, p 94–114.

82. Oton, J.; Sorzano, C. O.; Marabini, R.; Pereiro, E.; Carazo, J. M., Measurement of the modulation transfer function of an X-ray microscope based on multiple Fourier orders analysis of a Siemens star. Opt Express 2015, 23 (8), 9567–72.

83. Kremer, J. R.; Mastronarde, D. N.; McIntosh, J. R., Computer visualization of three-dimensional image data using IMOD. J Struct Biol 1996, 116 (1), 71–6.

84. Agulleiro, J. I.; Fernandez, J. J., Fast tomographic reconstruction on multicore computers. Bioinformatics 2011, 27 (4), 582–3.

85. Team, R. C. In R: A language and environment for statistical ## computing, R Foundation for Statistical Computing, 2020.

86. Goddard, T. D.; Huang, C. C.; Meng, E. C.; Pettersen, E. F.; Couch, G. S.; Morris, J. H.; Ferrin, T. E., UCSF ChimeraX: Meeting modern challenges in visualization and analysis. Protein Sci 2018, 27 (1), 14–25.

87. Frank, J.; Radermacher, M.; Penczek, P.; Zhu, J.; Li, Y.; Ladjadj, M.; Leith, A., SPIDER and WEB: processing and visualization of images in 3D electron microscopy and related fields. J Struct Biol 1996, 116 (1), 190–9.

88. Schneider, C. A.; Rasband, W. S.; Eliceiri, K. W., NIH Image to ImageJ: 25 years of image analysis. Nat Methods 2012, 9 (7), 671–5.

89. Benaglia, T.; Chauveau, D.; Hunter, D. R.; Young, D., mixtools: An R Package for Analyzing Finite Mixture Models. Journal of Statistical Software 2009, 32 (6).

